# Accelerating RepeatClassifier Based on Spark and Greedy Algorithm with Dynamic Upper Boundary

**DOI:** 10.1101/2021.06.03.446998

**Authors:** Kang Hu, Xingyu Liao, You Zou, Jianxin Wang

## Abstract

Transposable elements (TEs) represent quantitatively important components of genome sequences (e.g. 90% of the wheat genome), and play important roles in genome organization and evolution. The promotion of unsupervised annotation of transposable elements is of great significance. Classification is an important step in TE annotation, which summarize the information about the type or mechanism for the raw repetitive sequences. RepeatClassifier is a basic homology-based classification tool which compares the TE families to both the Repeat Protein Database (DB) and libraries of RepeatMasker. Unfortunately, RepeatClassifier is inefficient and takes a few days to classify the repetitive sequences of large genomes. Hence, we proposed Spark-based RepeatClassifier (SRC) which uses Greedy Algorithm with Dynamic Upper Boundary (GDUB) for data division and load balancing, and Spark to improve the parallelism of RepeatClassifier. Experimental results show that SRC can not only ensure the same level of accuracy as that of RepeatClassifier, but also achieve 42-88 times of acceleration compared to RepeatClassifier. At the same time, SRC shows excellent parallel performance when dealing with input datasets with unbalanced length distribution. SRC is publicly available at https://github.com/BioinformaticsCSU/SRC.

## 2 Introduction

Repetitive elements exist in most eukaryotes, and repetitive DNA sequences are fragments that appear multiple times in a genome. Repetitive DNA sequences can be divided into interspersed repetitive sequences and tandem repetitive sequences according to their structure. Interspersed repetitive sequences are mainly derived from transposable elements (TEs), which are highly identical and distributed separately in the genome(Treangen and Salzberg, 2012; Duan *et al*., 2019; Huang *et al*., 2012; Bourque *et al*., 2018; Guo *et al*., 2018). Tandem repeats are adjacent to each other and consist of satellites and simple repeats. Numerous studies have shown that the repetitive elements play an important roles in the evolution, inheritance, variation, gene expression, transcriptional regulation, chromosome construction, and physiological metabolism of living organisms(Lu *et al*., 1993; Kundu, 1999; Biscotti *et al*., 2015; Kaltenegger *et al*., 2018), and they are one of the principal causes of genomic instability(Lu *et al*., 2015).

TE annotation consists of three distinct steps: identification, classification, and masking (Feschotte and Pritham, 2007). Since the types and sequences of TEs are highly variable across species, Ab initio repeat identification is theoretically challenging and computationally intensive. Software packages like RECON (Bao and Eddy, 2002), RepeatScout (Price *et al*., 2005), Piler (Edgar and Myers, 2005), and ReAS (Li *et al*., 2005) have been designed to automate this process. Individually, none of these programs is able to generate a comprehensive, ‘masking-ready’ library of consensus repeats from an input genome sequence, but they produce a useful output representing the most abundant and homogeneous repeat families in a genome. The identification of repetitive sequences usually produces the raw TE consensus sequences, but does not provide information about the type or mechanism for transposition of the reconstructed repeat. The next step in TE annotation is to identify the diagnostic features of each consensus sequence, thereby inferring the biological classification of each repeat. RepeatModeler2 (Flynn *et al*., 2020), which together with other tools integrates them into one pipeline, make an attempt to classify the newly reconstructed consensus repeat, using sequence similarity to known repeats. Masking is the most straightforward as it consists of scanning the genome with sensitive algorithms for segments of the genome with significant similarity to one of several repeats precharacterized for the species and stored in a library of representative consensus sequences. The RepeatMasker software (http://www.repeatmasker.org/), which makes use of ‘manually-curated’ reference libraries of consensus sequences (Jurka *et al*., 2005), has become the gold standard for masking.

TEclass uses machine learning support vector machine (SVM) for classification based on oligomer frequencies(Abrusán *et al*., 2009). TEclass is a tool to classify unknown transposable elements into four main functional categories, which reflect their mode of transposition: DNA transposons, long terminal repeats (LTRs), long interspersed nuclear elements (LINEs) and short interspersed nuclear elements (SINEs). However, it cannot identify more detailed categories. REPCLASS uses three classification modules to classify repetitive sequence, including homology (HOM), structure (STR), and target site duplication (TSD)(Feschotte *et al*., 2009). RepeatModeler2 contains a basic homology-based classification module (RepeatClassifier) which compares the TE families generated by the various *de novo* tools to both the RepeatMasker Repeat Protein Database (DB) and to the RepeatMasker libraries (e.g., Dfam and/or RepBase). The Repeat Protein DB is a set of TE-derived coding sequences that covers a wide range of TE classes and organisms. As is often the case with a search against all known TE consensus sequences, there will be a high number of false positive or partial matches. RepeatClassifier uses a combination of score and overlap filters to produce a reduced set of high-confidence results. If there is a concordance in classification among the filtered results, RepeatClassifier will label the family using the RepeatMasker/Dfam classification system and adjust the orientation (if necessary). Remaining families are labeled Unknown if a call cannot be made. Classification is the only step that requires a database, and can be completed with only open-source Dfam if Repbase is not available. Unfortunately, The number of raw TE consensus repetitive sequences generated by various *de novo* tools may be huge, and the length of sequence may be particularly long. In this case, RepeatClassifier requires a extremely long running time, which seriously hinders the process of TE annotation in a genome sequence progress.

## 3 Methods

### 3.1 Parallel possibility analysis of RepeatClassifier

When classifying raw TE consensus repetitive sequences with RepeatClassfier, we need to go through the following steps (1)using *TRF* and *RepeatMasker* to scan for *low complexity* and *tandem* repeats. (2)comparing sequences with database of transposon proteins. (3)comparing sequences with ^*I\**^manually-curated^*I\**^ libraries of RepeatMasker. RepeatClassifier can be configured to use any of the comparison tools in *rmblast, abblast, wublast* for (2) (3) steps. No matter *TRF, RepeatMasker, rmblast, abblast* or *wublast*, they all have parallelization capabilities. Therefore, RepeatClassifier has the possibility of parallelism in theory.

### 3.2 Greedy Algorithm with Dynamic Upper Boundary (*GDUB*)

From the three steps of RepeatClassifier, the running time of the program should be influenced by the length of the repetitive sequence. To prove our conjecture, we make statistics on the change of running time with length. As shown in Fig. 3(A1 and A2), the running time showed a linearly correlated trend with the length of the sequence on the 9 datasets, including 7 species (**Supplementary Table S1**).

The length distribution of raw TE consensus repetitive sequences generated by *de novo* methods (**Supple mentary Section S1**) is unbalanced. As shown in Fig. 4(A1 to A9), the sequences are mainly concentrated below 1000bp in length, while there are also some ultra-long sequences concentrated above 3000bp. The unbalanced length distribution of sequences will inevitably lead to the unbalanced running time of each task in parallel mode. To make parallelism more efficient, we use Greedy algorithm with Dynamic Upper Boundary to make tasks more balanced. The pseudo code of GDUB is shown in **Supplementary Algorithm 5**.

### 3.3 Principle of Spark-based RepeatClassifier (*SRC*)

Resilient Distributed Datasets (*RDD*) are collection of various data items which are huge in size, that they have to be partitioned across various nodes. A partition in Spark is an atomic chunk of data (logical division of data) stored on a node in the cluster and it is basic unit of parallelism in Spark. Partitions are generated and distributed across different nodes automatically, which help parallelize distributed data processing with negligible network traffic for sending data between executors. The number of partitions determines the parallelism of Spark Program, we use GDUB algorithm to mark each sequence. The sequences in the same partition divided by GDUB will be marked with the same header. At the same time, the custom partition function used in Spark divide sequences with the same value (*header*%*partition_number*) into the same partition, which make the sum of sequence lengths in each partition as balanced as possible.

To ensure the accuracy of classification and improve the operating efficiency of the program, we use the Spark pipe method to invoke the original RepeatClassifier. This method can not only improve the parallel efficiency of the program, but also ensure the consistency of classified results. The processing principle of balanced allocation of repetitive sequences library generating by SRC is shown inFig. 2.

**Fig. 1.**
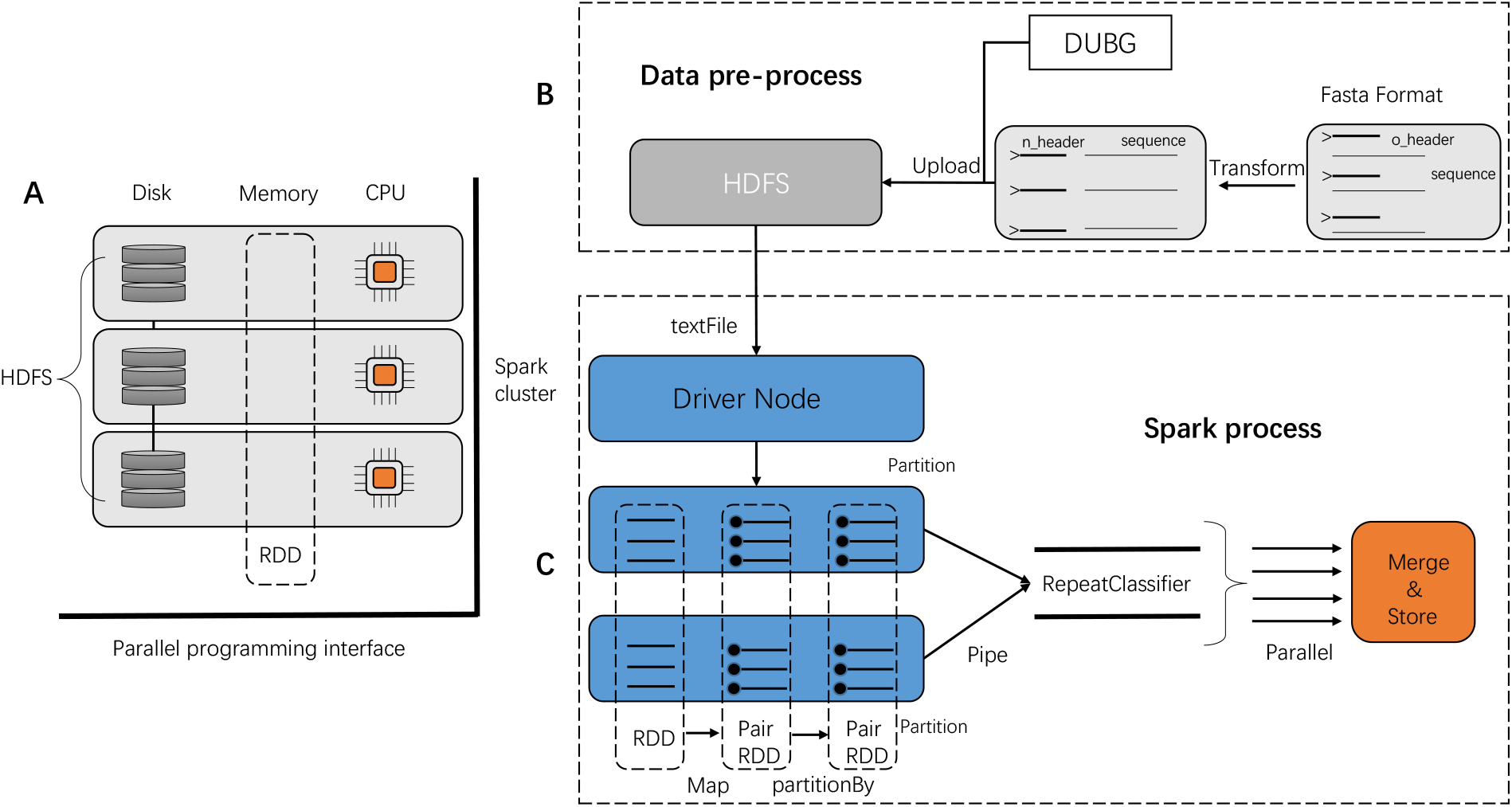
The sub-graph(A) shows the architecture of a Spark cluster and HDFS (Hadoop Distributed File System). The sub-graph(B) shows the process of data preprocess. The sub-graph(C) shows the process of Spark application. The dotted box in sub-graph(A) represents the resilient distributed datasets (*RDD*) of Spark, and the rectangles of grey color represent computing nodes in a cluster. Rectangles of light grey color in sub-graph(B) represent the unclassified repeat sequences files. Rectangles of blue color in sub-graph(C) represent computing nodes in the cluster, the dotted boxes represent RDDs in Spark and their transformation, a pipe formed by two thick black lines represents the use of RepeatClassifier program, and the rectangle of orange color represents the classified repeat sequences file.

**Fig. 2.**
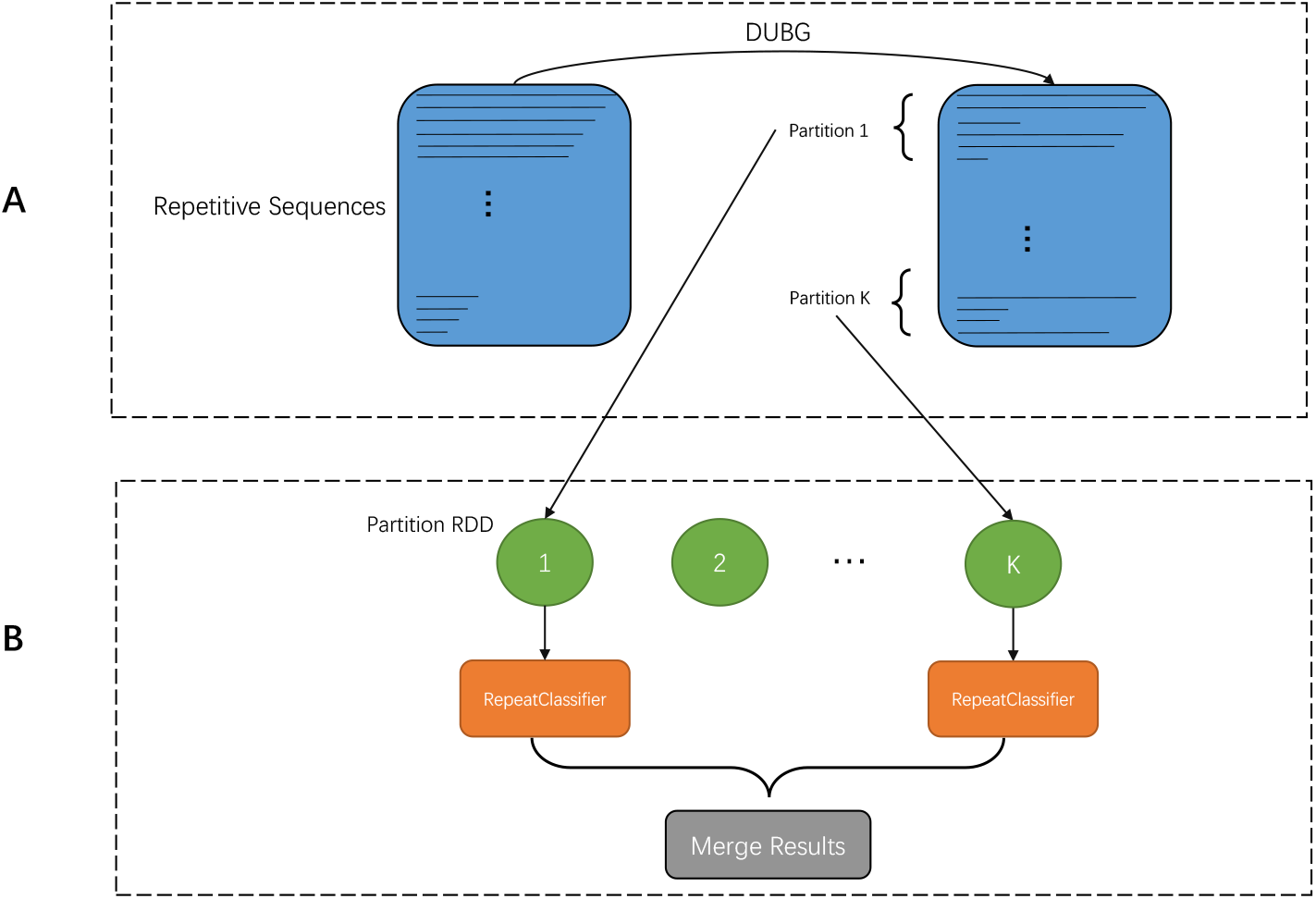
The processing principle of balanced allocation of repetitive sequences library. The sub-graph(A) shows the principle of generating partitions by GDUB algorithm. The sub-graph(B) shows the principle of the parallel computing model in SRC. Rectangles of blue color in Sub-graph(A) represent the unclassified repetitive sequence library (*fasta format*). Rectangles of green color in sub-graph(B) represent the corresponding partition in RDD of Spark, rectangles of orange color represent the applying of RepeatClassifier, and the rectangle of darkgrey color represent the final merged results.

## 4 Results

The hardware configuration of machines in all benchmarkings is shown in **Supplementary Fig S1**. The basic attributes of datasets are shown in **Supplementary Table S1**. All tools, commands and parameters are shown in **Supplementary Table S2**.

To study the relationship between the running time of RepeatClassifier and the length of input sequences, we extracted the first 10 longest sequences from original 9 datasets to form 9 smaller datasets. As shown in Fig. 3(A1 and A2), the running time shows a linear growth trend with the length of the sequence on all 9 datasets involving 7 species. To illustrate the characteristics of the input datasets, the length distribution of them are summarized. As shown in Fig. 4(A1 to A9), most of the sequences are concentrated in a short-length range (e.g., 0-400bp), but there are also some extremely long sequences existed (e.g., 495600bp).

**Fig. 3.**
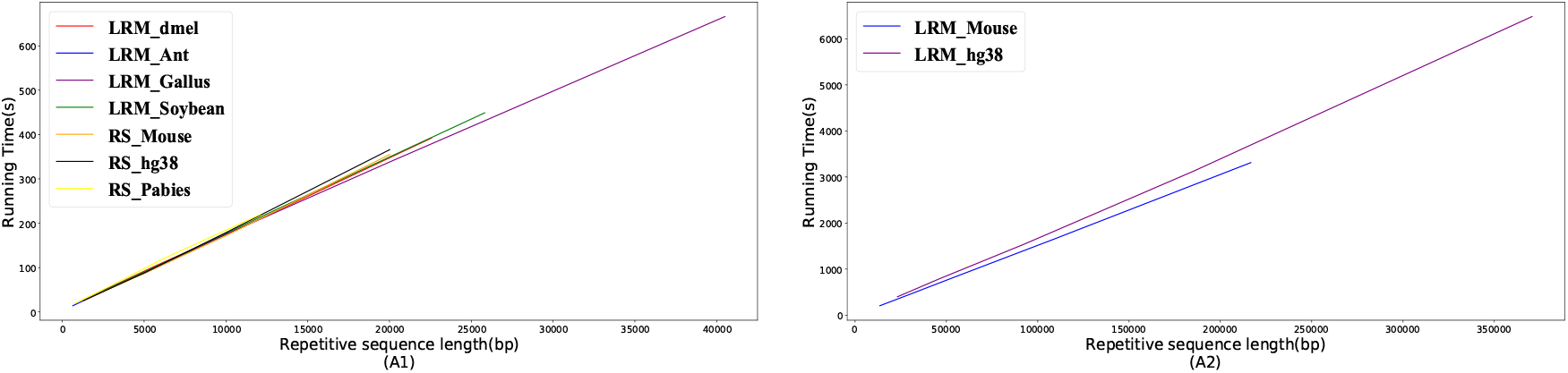
The relationship between the running time of RepeatClassifier and the length of sequences.

**Fig. 4.**
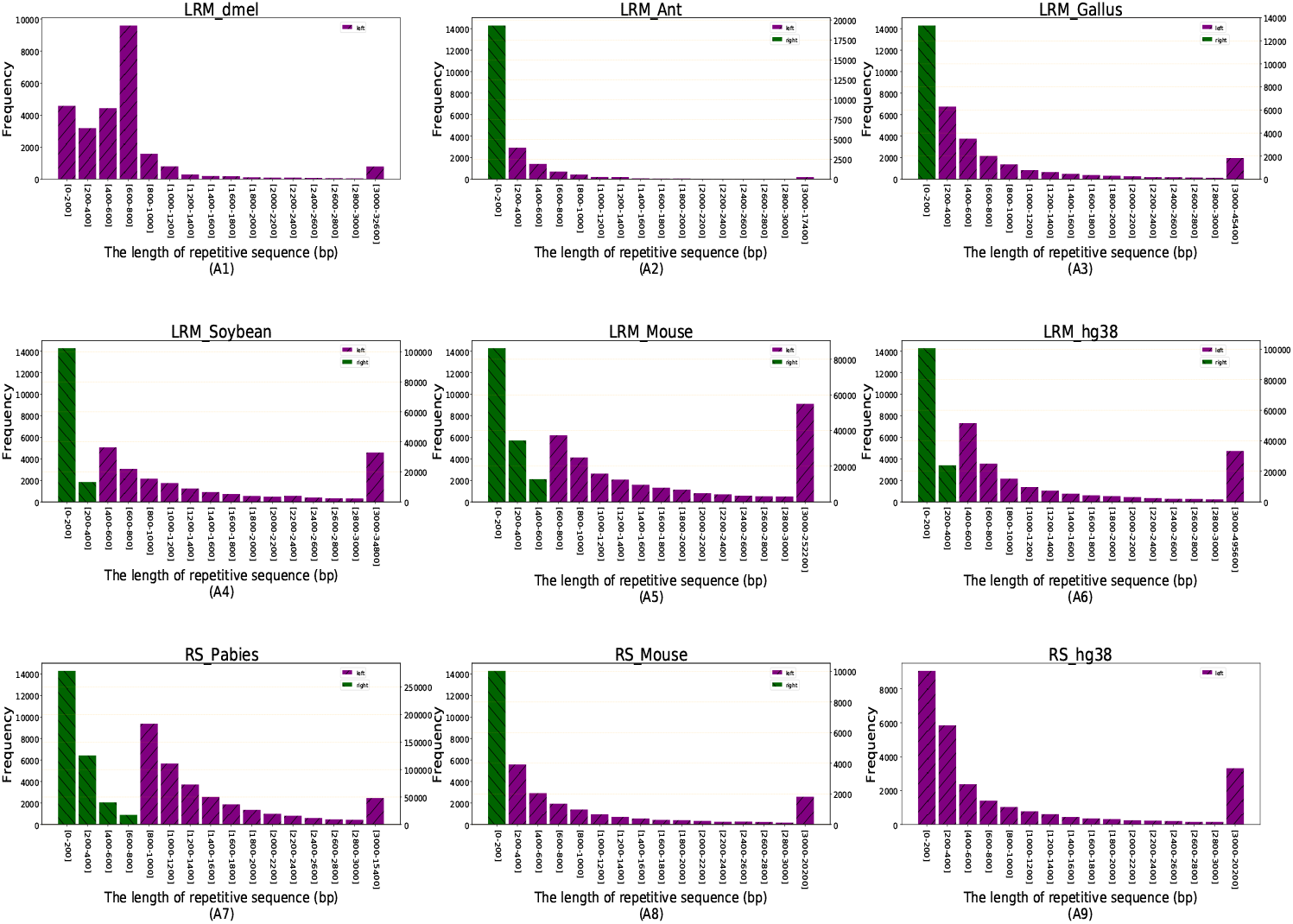
The length distribution of repetitive sequences. Note that two y-axes may exist in sub-graphs(A1-A9) since the frequency difference of the dataset is too large to be represented by only one y-axis. Purple bars correspond to the left y axis, which mean the frequency of repetitive sequence *<*= 10000, and green bars correspond to the right y axis, which mean the frequency of repetitive sequence *>* 10000.

Multiple partitioning methods are compared in **Supplementary Section S3**.**3**.**1**. As results, GDUB algorithm is the most task-balanced method, so it is chosen as the data partitioning method of SRC. To evaluate the performance of SRC, the comparison of SRC and RepeatClassifier is shown in Table 1. The Spark cluster is consisted of 5 nodes, and the detailed configuration of each node is shown in **Supplementary Fig S1**. Standalone mode is used in Spark and total executor cores are set to 240. At the same time, RepeatClassifier can only run on a single node, while SRC can run on a cluster. As shown in Table 1, the running time of SRC is only 1/88 to 1/42 of that of RepeatClassifier.

**Table 1.** Comparison of running time between SRC and RepeatClassifier

| Datasets | RepeatClassifier (min) | SRC (min) |
| --- | --- | --- |
| LRM_dmel | 592 | 9.8 |
| LRM_Ant | 195 | 4.6 |
| LRM_Gallus | 775 | 12 |
| LRM_Soybean | 1834 | 26 |
| LRM_Mouse | 3932 | 59 |
| LRM_hg38 | 3836 | 72 |
| RS_Pabies | 5053 | 57 |
| RS_Mouse | 940 | 13 |
| RS_hg38 | 1170 | 16 |

The changes of running time of SRC with the total executor cores are shown in Fig. 5(A1 to A9). In theory, the running time should be reduced by half when the total executor cores doubles, as shown by the red dotted line. However, the actual running time of SRC is shown by the green bar. The reason is that multiple threads are used in the process of RepeatClassifier, which causes the number of cores used by SRC reaching the threshold of the total number of cores in Spark cluster early. Detailed explanation is shown in **Supplementary Section S3**.**3**.**4**.

**Fig. 5.**
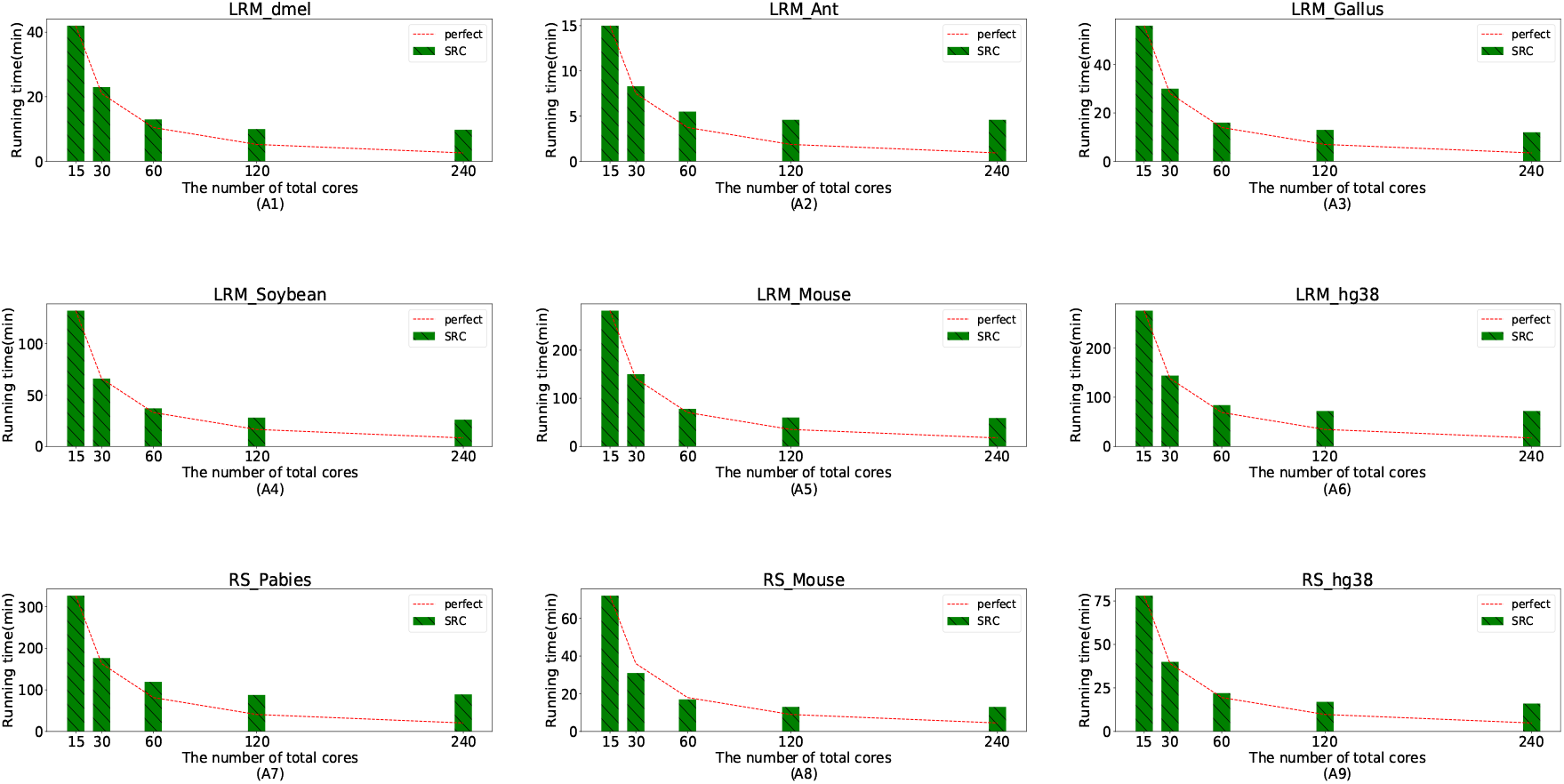
The running time of SRC changed with the total executor cores. The red dotted line in sub-graphs(A1-A9) represent the running time should be reduced by half when the total executor cores doubles theoretically and the green bar represent the actual running time of SRC.

## 5 Discussion

### 5.1 Consistency analysis of results

SRC is encapsulated on the basis of the original program of RepeatClassifier, and does not change its execution logic. In theory, the classified results should be exactly the same as that of RepeatClassifier, but there are slightly differences through real experiments. We find that the differences are caused by RepeatClassifier itself. For example, we randomly shuffle the input sequences of RepeatClassifier and get a slightly different result of classification. This problem can be resolved with the upgrade of RepeatClassifier. To prove that SRC and RepeatClassifier have the same level of accuracy, RepeatMasker is used to cover ^*I\**^manually-curated^*I\**^ reference libraries extracted from RepeatMasker using *queryRepeatDatabase*.*pl*. The detailed experimental results are shown in **Supplementary Tables S4 to S12**. Although the results of SRC and RepeatClassifier are slightly different, the total coverage is basically the same, which is shown in Fig. 6. In addition to covering libraries, Minimap2 and BWA are used to cover genome references. The detailed results are shown in **Supplementary Table S13 to S21**. From the results shown in **Supplementary Tables S4 to S21**, we can conclude that SRC has the same level of accuracy as that of RepeatClassifier.

**Fig. 6.**
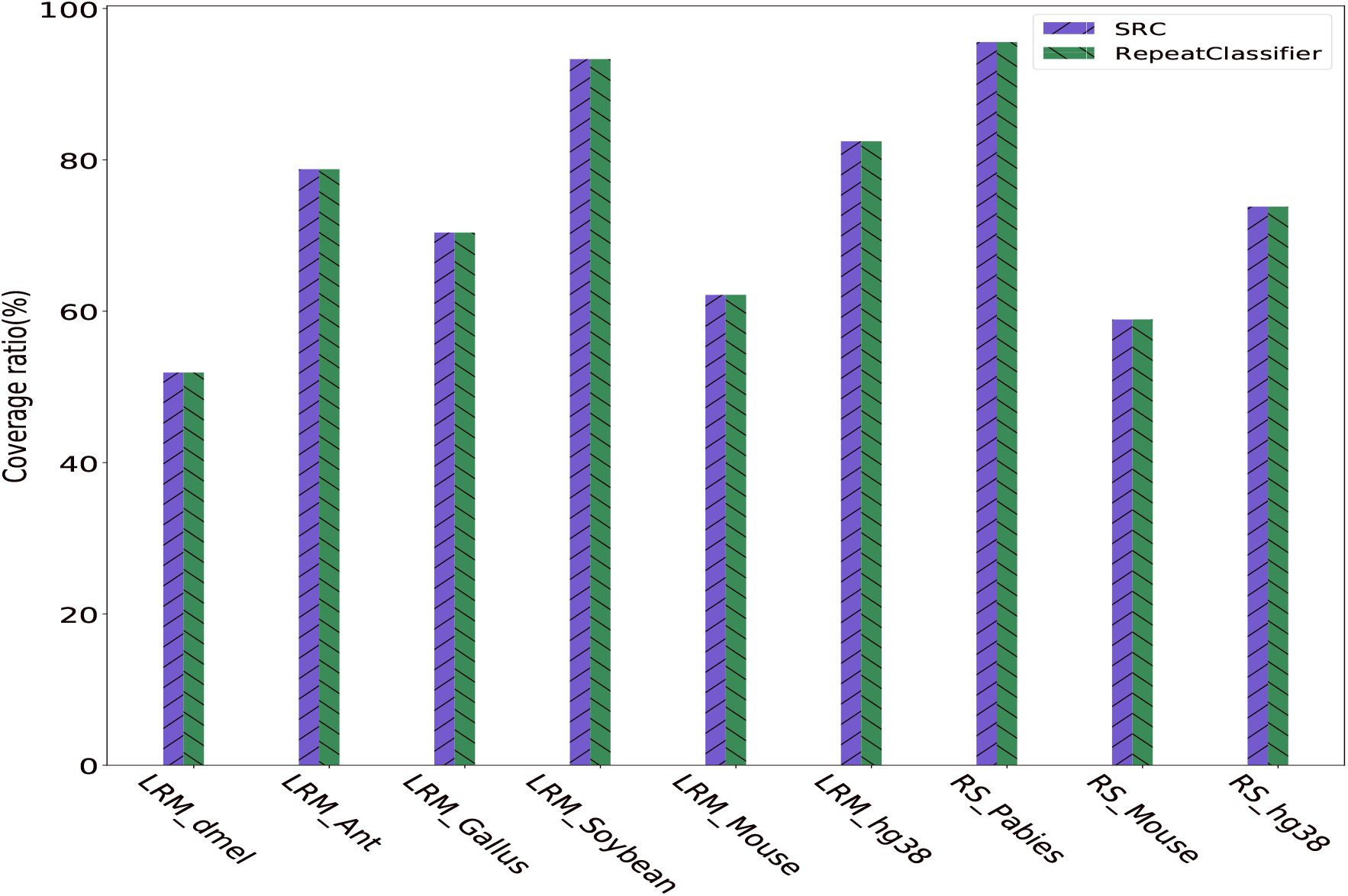
The total proportion classification of detection results of SRC and RepeatClassifier covering RepeatMasker libraries.

### 5.2 Future work

The process of TE annotation in a genome sequence consists of three distinct steps: identification, classification, and masking. RepeatClassifier is aimed at the classification of raw repetitive sequences. In this paper, we propose SRC to accelerate the classification of TE annotation, but we do not involve the other two steps. In the future, we can study how to accelerate the the other two steps of TE annotation.

## 6 Conclusion

In this paper, we proposed Spark-based RepeatClassifier (SRC) for TE classification, which not only keep the same execution logic as that of RepeatClassifier, but also achieve a higher running efficiency (42-88 times) compared to RepeatClassifier. At the same time, a modular interface is provided to facilitate the subsequent upgrade and optimization of RepeatClassifier.

## 7 ACKNOWLEDGEMENTS

This work has been supported by the National Natural Science Foundation of China under Grant:No.61772557, No.61732009 and No.62002388, Hunan Provincial Science and technology Program (No. 2018wk4001), 111 Project (No. B18059), Fundamental Research Funds for the Central Universities of Central South University (2021zzts0208).

