## Supplementary File for "Accelerating RepeatClassifier Based on Spark and Greedy Algorithm with Dynamic Upper Boundary"

### 1 Classification of existing repeat detection methods

As shown in Fig. S1, there are three categories of detection methods for repetitive regions in genome, including homology-based, structure-based and *de novo* methods. RepeatMasker is a typical method in homology based methods, which searches homology databases to find and mask the repetitive sequences. The structure-based identification methods take advantage of the prior information about the structure features in sequences, and identify the repetitive sequences based on heuristic algorithms(Jiang, 2013). The *de novo* methods are more flexible than the other two methods since they do not require repetitive structure or prior information about similarity to known repetitions(Shi and Liang, 2019). The *de novo* methods can be further divided into three categories: the multiple sequence alignment-based methods, the *k-mer* and space seed extension-based methods, and sequence assembly and similarity network-based methods. The first one relies on the multiple sequence alignment to identify repeats, including Repeat Pattern Toolkit(Agarwal *et al.*, 1994), RECON(Bao and Eddy, 2002), PILER(Edgar and Myers, 2005) and LTRdigest(Nicolas *et al.*, 2016). The second category relies on *k-mer* and space seed extension strategies to identify repetitive sequences, including ReAS(Li *et al.*, 2005), RepeatScout(Price *et al.*, 2005), RepeatFinder(Saha *et al.*, 2008), EDTA(Ou *et al.*, 2019) and Generic Repeat Finder (GRF)(Shi and Liang, 2019). The methods in the third category rely on sequences assembly and similarity network to identify repeats, including RepARK(Koch *et al.*, 2014), REPdenovo(Chu *et al.*, 2016) and RepLong(Guo *et al.*, 2018).

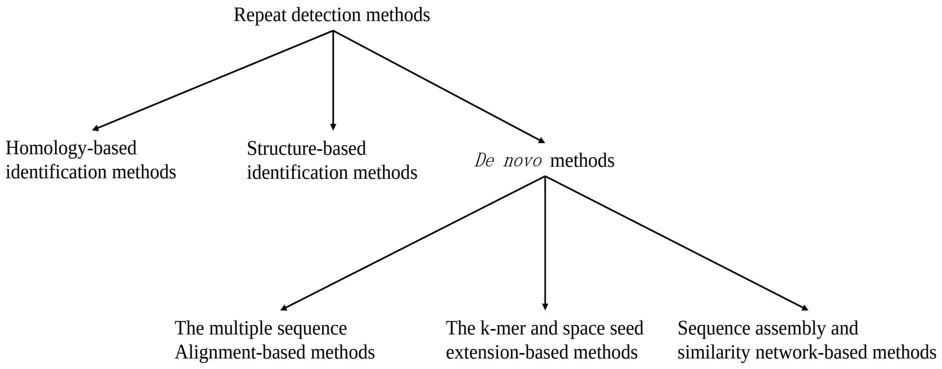


**Fig. S1.** The classification of detection methods.

### 2 Methods

#### 2.1 Multi-Way Number Partitioning

The goal of this study is to allocate tasks to different executors on Spark and make the amount of tasks on each executor is as close as possible, which is a multiprocessor scheduling problem(Hou *et al.*, 1994; Grigoreva, 2020). Multiprocessor scheduling problem is defined as follows: The input set $S$ is composed of $n$ positive integers, which corresponds to the running time of a set of $n$ jobs. The number of subsets $k$ corresponds to a number of identical machines, such as processor cores that execute in parallel. The goal of multiprocessor scheduling is to assign each of jobs to one of the $k$ machines while minimizing the makespan, or the time required to complete all jobs in the schedule. Minimizing the makespan is equivalent to minimizing the time to complete all jobs on the machine with the longest running time. Multiprocessor scheduling problem can be equivalent to the problem of multi-way number partitioning which is described in detail as follows.

Let $S = (\alpha_1, \alpha_2, ..., \alpha_n)$ be a list of $n$ positive integers, $q_i$ be the sum of the numbers in the $i$-th subset. The number partitioning problem is the task of partitioning $S$ into $k$ subsets $S_1, S_2, ..., S_k$ so that the sum of the numbers in different subsets are as nearly equal as possible. For instance, if $S = (1, 1, 2, 3, 4, 5)$ and $k = 2$, we can consider the following partition $(1, 1, 2, 4)$ and $(3, 5)$. The numbers in each subset add up to 8, so this is a completely balanced partition. There are three objective functions are proposed for this problem(Ahmadypour and Gohari, 2020).

1) [Min-Difference objective function] Minimize the difference between the largest and smallest subset sums, i.e., minimize $(\max_{q_i} - \min_{q_i})(1 \le i \le k)$.
2) [Min-Max objective function] Minimize the largest subset sum, i.e., minimize $\max_{q_i}(1 \le i \le k)$.
3) [Max-Min objective function] Maximize the smallest subset sum, i.e., maximize $\min_{q_i}(1 \le i \le k)$.

While these objective functions are equivalent when $k = 2$, neither of them is equivalent to the other for $k > 2$. The objective function of our task is Min-Max objective function.

**2.1.1 Direct Partitioning Algorithm** is the simplest strategy of partitioning. The sequences are sorted by length and evenly divided into each partition. The sequences in each partition keep the original order, and the core processing steps of this partitioning strategy are described by *Algorithm 1.* It divides the sequences into partitions evenly, regardless of the length distribution of the repetitive sequences, which will inevitably lead to extremely unbalanced load of tasks. Consider the input set $S = \{18, 17, 12, 11, 8, 2\}$ and the number of partition $k = 2$. The size of $S$ is 6 and $k$ equals to 2, so the size of each partition equals to $\frac{6}{2} = 3$. Since $S$ is already in descending order, the final partition result is $< \{18, 17, 12\}, \{11, 8, 2\} >$. The upper bound of cost is $\max < \sum S_1, \sum S_2 > = \max < 47, 21 > = 47$.

**Algorithm 1** Direct partitioning algorithm

**procedure** DIRECT($S, k$) ▷ $S$ represents the set used to store the numbers, $k$ be the number of partitions
  **Let $\prod$ be the set used to store partitions of numbers**
  **Let $p$ be the index of partition**
  **Let $avg$ be the average of the sum of all numbers**
  **Let $i$ be the index to record current traversal location**
  $\prod \leftarrow [[] \overset{k}{\ldots} []]$
  $avg \leftarrow sum(S)/k$
  $p \leftarrow 0$
  $i \leftarrow 0$
  $S.sortDescend()$
  **while** $S \neq []$ **do**
    $w \leftarrow S.popFirst()$
    **if** $i\%avg == 0$ **and** $i \neq 0$ **then**
      $p = (p + 1)\%k$
    **end if**
    $\prod[p].append(w)$
    **increment** $i$
  **end while**
  **return** $\prod$
**end procedure**

**2.1.2 Paired End Traversal Algorithm (PET)** is a heuristic algorithm based on the strategy of dividing the long sequence and the short sequence into the same partition, and ensure the total sequence length of each partition as balanced as possible, which has obvious advantages compared with the direct partitioning method. For a repetitive sequences set $R$ of size $|R| = n$, *traversal_direction* is denoted as the direction of sequences traversal, 1 or 0 represents the direction is forward or reverse. $i$ and $j$ are denoted as the position of forward and reverse traversal, respectively. *PET* starts traversing from the head of the repetitive sequences ($i \leftarrow 0$, $j \leftarrow n\text{-}1$). *node_index* are used to represent the increasing marking-index and *partition_num* is denoted as the partition number of Spark jobs. While the number of traversed sequences reaches the number of partitions, change the direction of sequences traversal, and continue from last traverse position in the same direction. The details of PET are described by *Algorithm 2.* Consider the input set $S = \{18, 17, 12, 11, 8, 2\}$. The following are the steps the PET algorithm takes to compute an upper bound on the cost of a two-way partition:(1)The direction of sequences traversal is forward. Pop the maximum number from $S$, and push it into the first partition. The state of partition is $< \{18\}, \{\} >$. (2)Pop the maximum number remaining in $S$, and push it into the second partition. The state of partition is $< \{18\}, \{17\} >$. (3)Since the count of pushed number reaches an integer multiple of the number of partition, the direction of sequences traversal is changed into reverse. Pop the minimum number from $S$, and push it into the first partition. The state of partition is $< \{18, 2\}, \{17\} >$. (4)Pop the minimum number remaining in $S$, and push it into the first partition. The state of partition is $< \{18, 2\}, \{17, 8\} >$. (5)Since the count of pushed number reaches an integer multiple of the number of partition, the direction of sequences traversal is changed into forward.... (6)The final result of partition is $< \{18, 2, 12\}, \{17, 8, 11\} >$. The upper bound of cost is $\max < \sum S_1, \sum S_2 > = \max < 32, 36 > = 36$.

**2.1.3 Greedy Algorithm** makes the locally optimal choice at each stage and with hope of finding a global optimum. For a input set $S$ of size $|S| = n$, the greedy algorithm first sorts the input set $S$ into monotonically decreasing order. It then considers the integers one at a time and places them into the subset with the smallest cumulative sum. If there are several subsets with the smallest cumulative sum, one of the subsets is chosen arbitrarily. The time complexity of the algorithm is $O(nlogn)$ and the space complexity is $O(n)$(Barat, 2017). The partition costs are within 4/3 of optimal(Kellerer *et al.*, 2004). The greedy algorithm is optimal for n $\leq$ 4(Korf, 2011). The details of greedy algorithm are described by *Algorithm 3.* Consider the input set $S = \{18, 17, 12, 11, 8, 2\}$ and the number of partitions equals to 2. The following are the steps the greedy algorithm takes to compute an upper bound on the cost of a two-way partition:(1)Pop the two largest numbers and push them into different partitions. The state of partition is $< \{18\}, \{17\} >$. (2)Pop the largest number remaining in $S$, and push it into the partition with the smallest sum. The state of partition is $< \{18\}, \{17, 12\} >$. (3)Iterate until all numbers are pushed into the partitions. The final state of partition is $< \{18, 11, 8\}, \{17, 12, 2\} >$. The upper bound of cost is $\max < \sum S_1, \sum S_2 > = \max < 37, 31 > = 37$.

**2.1.4 The Karmarkar-Karp Set Differencing Algorithm (KK)** provides an alternative to the greedy algorithm (Karmarkar and Karp, 1982). Like the greedy algorithm, KK begins by sorting $S$ into decreasing order. Then, it iteratively replaces the largest two integers of $S$ with their difference. This is equivalent to placing the two integers into separate subsets without specifying which integer goes into which subset. KK continues in this manner, replacing the two largest integers with their difference until there is only one integer left, which is the difference between the sums of the final sets, $|sum(S_1) - sum(S_2)|$. Given this difference, the larger subset sum is $\frac{sum(S)+|sum(S_1)sum(S_2)|}{2}$. The details of the KK algorithm are described by *Algorithm 4.* Consider the input set $S = \{18, 17, 12, 11, 8, 2\}$ and the number of partitions equals to 2. The 18 and 17

**Algorithm 2** PET partitioning algorithm

```
procedure PET(S, k)        ▷ S represents the set used to store the numbers, k be the number of partitions
    Let ∏ be the set used to store partitions of numbers
    Let d be the flag used to record the direction of sequences traversal, 1 represents forward and 0 represents reverse
    Let p be the index to record the count of traversal
    Let i be the index used to record the traversal position from the head
    Let j be the index used to record the traversal position from the tail
    ∏ ← [[] ...k []]
    d ← 1
    p ← 0
    i ← 0
    j ← len(S) − 1
    S.sortDescend()
    while S ≠ [] do
        w ← S.popFirst()
        ∏[p%k].append(w)
        if d then
            increment i        ▷ go forward
        else
            decrement j        ▷ go reverse
        end if
        increment p
        if p % k == 0 then
            d = Bool(1 − d)        ▷ Change the direction of sequences traversal
        end if
    end while
    return ∏
end procedure
```

**Algorithm 3** Greedy partitioning algorithm

```
procedure Greedy(S, k)        ▷ S represents the set used to store the numbers, k be the number of partitions
    Let ∏ be the set used to store partitions of numbers
    Let ∑ be the set used to store the sum of numbers in each partition
    ∏ ← [[] ...k []]
    ∑ ← [0 ...k 0]
    S.sortDescend()
    while S ≠ [] do
        w ← S.popFirst()        ▷ Largest remaining number
        p_min ← argmin(∑)        ▷ Index of lightest part
        ∏[p_min].append(w)
        ∑[p_min] ← ∑[p_min] + w
    end while
    return ∏
end procedure
```

are replaced by their difference of 1, which is inserted into the remaining list, resulting in $\{12, 11, 8, 2, 1\}$. Next, the 12 and 11 are replaced by their difference of 1, resulting in $\{8, 2, 1, 1\}$. The 8 and 2 are replaced by 6, resulting in $\{6, 1, 1\}$. The 6 and 1 are replaced with 5, resulting in $\{5, 1\}$. Their difference results in the final subset difference of 4. The cost is $\frac{sum(S)+difference}{2} = \frac{68+4}{2} = 36$, corresponding to the partition: $< \{18, 12, 2\}, \{17, 11, 8\} >$, with subset sums of 32 and 36, respectively. To construct the actual subsets, a graph is created with a node for each integer. Whenever two integers are replaced by their difference, an edge between the two nodes is added. The new value is represented by the node with the larger value. Finally, the resulting graph is two-colored to construct the two subsets.

**Algorithm 4** Karmarkar-Karp partitioning algorithm

**procedure** KK$(S, k)$ ▷ $S$ represents the set used to store the numbers, $k$ be the number of partitions
    **Let $L_i$ be the $i$th element of the list L**
    $n \leftarrow length(S)$
    $S.sortDescend()$
    $M \leftarrow [[S_1, 0 \overset{k-1}{\dots} 0] \overset{n}{\dots} [S_n, 0 \overset{k-1}{\dots} 0]]$ ▷ Initialization: transform numbers into k-tuples
    $AmB, norm \leftarrow ([], [])$
    **while** $length(M) \geq 2$ **do**
        $A, B \leftarrow pop2First(M)$
        $AmB.append([[A_1, B_k] \overset{k}{\dots} [A_k, B_1]])$ ▷ Save the old elements
        $E \leftarrow [(A_1 + B_k) \overset{k}{\dots} (A_k + B_1)]$ ▷ Compute the new element
        $e_{min} \leftarrow \min(E)$ ▷ Normalization factor
        $norms.append(m)$ ▷ Save the normalization factor
        $E \leftarrow [E_1 - e_{min} \overset{k}{\dots} E_k - e_{min}]$ ▷ Normalize
        $E.sortDescend()$
        $M.InsertInOrderedMatrix(E)$ ▷ So that $M_1[1] \geq M_2[1] \geq ...$
    **end while**
    $\prod \leftarrow M_1$
    **while** $AmB \neq []$ **do**
        $E, m \leftarrow (AmB.popLast(), norms.popLast())$
        $P \leftarrow [1 \overset{k}{\dots} k]$
        **for** `[a,b]` $\in$ `E` **do**
            $p \leftarrow \prod(a + b - m)$ ▷ Index of the part (a+b-m) belongs to
            $P.remove(p)$
            $\prod_p .remove(a + b - m)$
            $\prod_p .append(a)$
            $\prod_p .append(b)$
        **end for**
    **end while**
    **return** $\prod$
**end procedure**

##### 2.1.5 Greedy Algorithm with Dynamic Upper Boundary (GDUB)

**Greedy Algorithm with Dynamic Upper Boundary (GDUB)** uses a dynamically increasing boundary to improve the shortcomings of the greedy algorithm. Like the greedy algorithm, GDUB algorithm begins by sorting $S$ into decreasing order and calculate an ideal value which is equal to the sum of the elements divided by the number of partitions rounded to the nearest integer. Then a dynamic boundary is generated based on the ideal value. According to this dynamic boundary, each value in $S$ will be divided into a suitable partition. If a value is not put into the appropriate partition in one iteration, the upper bound of the boundary will be increased before the next iteration to make the value be put into the appropriate partition as much as possible. GDUB algorithm has an overall time complexity of $O(n * k)$, $n$ being the number of elements and $k$ being the number of partitions. The details of the GDUB algorithm are described by *Algorithm 5*. A simple approximation example would be, consider the input set $S = \{18, 17, 12, 11, 8, 2\}$ and the number of partitions equals to 2. The ideal value is $\frac{sum(S)}{2} = \frac{68}{2} = 34$. The 8 and 17 are pushed into two partitions in turn, and the partition status is $< \{18\}, \{17\} >$. Trying to push 12 into the first partition, $sum(\{18, 12\}) = 30 < 34$, the partition status is $< \{18, 12\}, \{17\} >$. Trying to push 11 into the first partition, $sum(\{18, 12, 11\}) = 41 > 34$, which exceeds the ideal value. Trying to push 11 into the second partition, $sum(\{17, 11\}) = 28 < 34$, the partition status is $< \{18, 12\}, \{17, 11\} >$. Trying to push 8 into the second partition, $sum(\{17, 11, 8\}) = 36 > 34$. Since the last partition has been reached, the upper bound needs to be expanded to 35. There is still no appropriate partition to push 8. Expanding the upper bound to 36, and the second partition is appropriate, $sum(\{17, 11, 8\}) = 36 \geq 36$. The partition status is $< \{18, 12\}, \{17, 11, 8\} >$. In the same way, 2 can be pushed into the first partition, and the partition status is $< \{18, 12, 2\}, \{17, 11, 8\} >$.

#### 2.2 Process of SRC

After finishing the data preprocessing, the sequences can be input into the process of SRC. The process of SRC consists of 6 steps: (1)read file from HDFS (2)convert RDD to Paired RDD (3)partition RDD by custom partition function (4)add prefix for each partition (5)invoke RepeatClassifier program for each partition (6)merge all classified partition. The process of SRC can be abstracted into a DAG graph including two stages. As shown in Fig. S2, the first stage is for data reading and conversion, and the second stage is for custom partitioning of data and calling RepeatClassifier for calculation.

**Algorithm 5** Greedy algorithm with Dynamic Upper Boundary

```
procedure GDUB(S, k)    ▷ S represents the set used to store the numbers, k be the number of partitions
    Let ∏ be the set used to store partitions of numbers
    ∏ ← [[] …k… []]
    b ← sum(S)/k
    S.sortDescend()
    i ← 0
    while i < k do
        w ← S.popFirst()
        ∏[i].append(w)
        increment i
    end while
    while S ≠ [] do
        w ← S.popFirst()
        assigned = False
        p = 0
        while not assigned do
            if sum(∏[p]) + w <= b then
                ∏[p].append(w)
                assigned = True
            else
                if p == len(∏) − 1 then
                    p = 0
                    increment b
                else
                    increment p
                end if
            end if
        end while
    end while
    return ∏
end procedure
```

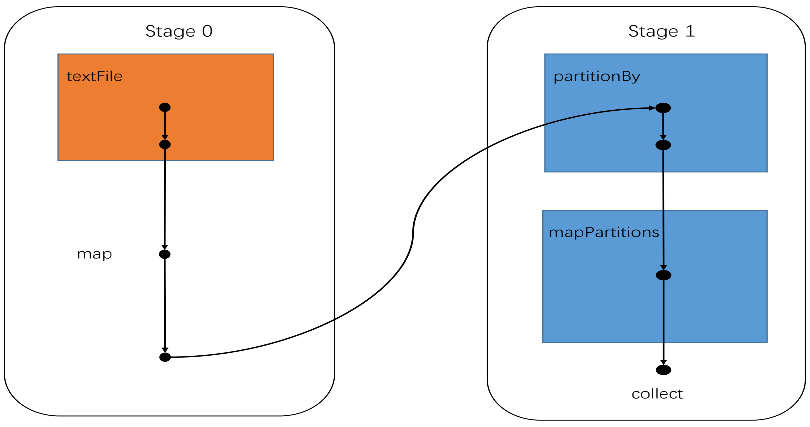


**Fig. S2.** The DAG graph of SRC.

### 3 Results

#### 3.1 Benchmarking datasets and configurations of tools

All experimental datasets are mainly divided into two parts: the repetitive sequences found by LongRepMarker (https://github.com/BioinformaticsCSU/LongRepMarker) and the repetitive sequences found by RepeatScout (http://bix.ucsd.edu/repeatscout/), which involving 7 species: Drosophila melanogaster, Acromyrmex echinatior, Gallus gallus bankiva, Glycine max, Mus musculus, Homo sapiens and Picea abies. The details of datasets are shown in Table S1 and the detailed commands of all benchmarking tools are shown in Table S2.

**Table S1. All species datasets used in benchmarkings**

| Tools | Species | abbreviation | Genome size (MB) | Sequences size (Mbp) | Number (Thousand) | Max length (kb) | Min length (bp) |
|---|---|---|---|---|---|---|---|
| LongRepMarker | Drosophila melanogaster | LRM_dmel | ~165 | 19 | 26.157 | 32.483 | 49 |
| | Acromyrmex echinatior | LRM_Ant | ~283 | 6.4 | 25.685 | 17.329 | 49 |
| | Gallus gallus bankiva | LRM_Gallus | ~1126 | 28 | 32.986 | 45.336 | 49 |
| | Glycine max | LRM_Soybean | ~934 | 60 | 137.711 | 34.741 | 49 |
| | Mus musculus | LRM_Mouse | ~2764 | 136 | 164.562 | 252.066 | 49 |
| | Homo sapiens | LRM_hg38 | ~3174 | 131 | 148.244 | 495.458 | 49 |
| RepeatScout | Picea abies | RS_Pabies | ~13312 | 138.28 | 492.858 | 15.309 | 50 |
| | Mus musculus | RS_Mouse | ~2764 | 29.54 | 28.761 | 20.016 | 50 |
| | Homo sapiens | RS_hg38 | ~3174 | 35.99 | 26.540 | 20.016 | 50 |

Note that the abbreviation of dataset corresponds to manuscript Fig.3.

#### 3.2 Hardware Configuration

All benchmarkings are done on computers with 48 cores (Intel(R) Xeon(R) Gold 6248R CPU @ 3.00GHz) and the memory of 192GB. The hardware configuration is shown in Fig. S3.

**Table S2. Details of Benchmarking tools**

| Tools | Benchmarking tools | | |
|---|---|---|---|
| | Version | Requirements | Running commands and parameters configuration |
| RepeatMasker | v4.0.7 | Perl 5.8.0 or higher; Python 3.0; h5py HDF5 library for Python; TRF 4.09 or higher; A search engine (crossmatch/ rmblast/nhmmer/abblast/wublast); Dfam and/or RepBase libraries. | queryRepeatDatabase.pl -species "taxon" > repeatmasker.taxon.fa.<br><br>RepeatMasker -parallel threads -lib custom_library -x -html -g -dir output_dir repeatmasker_library. |
| RepeatClassifier | v2.0.1 | Perl; RepeatMasker; RMBlast. | RepeatClassifier [-options] -consensi <repeat model file> [-stockholm <stockholm file>] [-engine <abblast\|ncbi>]. |
| Minimap2 | v2.17-r954-dirty | GNU g++ 4.0 | minimap2 -d index_path target.fa. minimap2 -a -t threads_num index_path query.fa. |
| BWA | 0.7.17-r1188 | GNU g++ 4.0 | bwa index -a bwtsw target.fa. bwa mem -t threads_num -x intractg -a target.fa query.fa. |
| RepeatScout | 1.0.5 | GNU g++ 4.0; Perl 5.5.0 or higher; TRF; | build_lmer_table -sequence reference -freq freq_path. RepeatScout -sequence reference -output output_dir -freq freq_path. |

| CPU | | RAM | | DISK | |
|---|---|---|---|---|---|
| Parameters | Value | Parameters | Value | Parameters | Value |
| Architecture | X86_64 | Total Mem | 187G | /dev/mapper/centos-root | 190G |
| CPU op-mode(s) | 32-bit, 64-bit | Total Swap | 31G | devtmpfs | 94G |
| Physical CPU | 2 | Buff/Cache | 3G | tmpfs | 94G |
| Processors | 48 | | | tmpfs | 94G |
| CPU cores | 24 | | | tmpfs | 94G |
| Model name | Intel(R) Xeon(R) Gold 6248R CPU @ 3.00GHz | | | /dev/sda2 | 1014M |
| CPU MHz | 3599.121 | | | /dev/sda1 | 200M |
| Virtualization | VT-x | | | 10.10.10.201@o2ib:10.10.10.202@o2ib:/ddnes | 2.1P |
| L1d cache | 32K | Operation System | | | |
| L1i cache | 32K | Parameters | Value | | |
| L2 cache | 1024K | Version | CentOS Linux release 7.5.1804 (Core) | | |
| L3 cache | 36608K | Kernel | Linux node1017 3.10.0-862.el7.x86_64 #1 SMP Fri Apr 20 16:44:24 UTC 2018 x86_64 x86_64 x86_64 GNU/Linux | | |

**Fig. S3.** Hardware Configuration.

### 3.3 Details of experiment results

**3.3.1 Comparison of partitioning methods** To compare the performance of each partition method, sufficient experiments are conducted on Direct, PET, Greedy, KK, and GDUB based on benchmarking datasets. Since the running time of RepeatClassifier is linearly correlated to the length of the sequence (Manuscript Figs. 4), the length of the sequence is used to represent the running time. The input data is a collection of sequence lengths and the number of partitions in the experiment is set to 240. As shown in Table S3, Direct, PET and KK algorithms are fast, while Greedy and GDUB consume longer running time. However, Greedy and GDUB algorithms are superior to the other three algorithms in terms of partition balance. To prove that the GDUB algorithm is the most suitable partitioning mehod for SRC, RS_Pabies dataset with the longest running time of GDUB is reused. As shown in Fig. S4, SRC-GDUB spends more time on partitioning, but its whole running time is still less than that of SRC-PET. Hence, GDUB is chosen as the partition algorithm of SRC.

**3.3.2 Task balance of Spark application** SRC-GDUB and SRC-Direct represent the SRC program using the GDUB and Direct data partitioning method. As shown in Fig. S5(A1), the running time of SRC-GDUB is only 1/21 to 1/3 of that of SRC-Direct method. To illustrate the effect of load balancing, we configure Spark history to view the running time of each task in Spark application. As shown in Fig. S6, the running time of tasks in direct data partitioning method is extremely unbalanced, with a small number of tasks occupying the main running time. On the contrary, the running time of tasks in GDUB method consumes almost consistent running time.

**3.3.3 Memory usage** As shown in Fig. S5(A2), the peak memory usage of RepeatClassifier varies greatly with different datasets, while that of SRC keeps relatively stable. As shown in Fig. S5(B1 to B9), the memory usage of all nodes in the cluster has a similar trend of change which rising rapidly and stabilizing, then falling rapidly, and maintaining a steady state until the end on the most datasets except for RS_Pabies dataset. We find there are extremely high frequency short sequences (e.g., 0-800 bp) shown in Fig. S7(A1). After many short sequences are filtered out, the sequences length distribution of the dataset is shown in Fig. S7(A2) and the memory usage of the cluster has a tendency to drop sharply, so it can be concluded that the slow down trend of the memory usage of the cluster is caused by too many high-frequency short sequences. Since the master node needs to maintain task scheduling and other tasks, it consumes more memory than other nodes.

**Table S3. Comparison of partitioning methods**

| Datasets | | Direct | PET | Karmarkar-Karp | Greedy | GDUB |
|---|---|---|---|---|---|---|
| LRM_dmel | time | **0.003766** | 0.004940 | 0.004766 | 3.370448 | 2.685078 |
| | max | 1071534 | 107014 | 107518 | 80017 | **80006** |
| | min | 6462 | 75119 | 75173 | 79967 | 79957 |
| | max-min | 1065072 | 31895 | 32345 | 50 | **49** |
| LRM_Ant | time | **0.003419** | 0.004809 | 0.005195 | 2.860052 | 2.590868 |
| | max | 624291 | 40258 | 40209 | **25935** | 25953 |
| | min | 5251 | 22986 | 22987 | 25886 | 25907 |
| | max-min | 619040 | 17272 | 17222 | 49 | **46** |
| LRM_Gallus | time | **0.004505** | 0.006501 | 0.005974 | 4.601177 | 4.007600 |
| | max | 3340747 | 151699 | 151747 | **119671** | 119698 |
| | min | 6842 | 106648 | 106566 | 119622 | 119650 |
| | max-min | 3333905 | 45051 | 45181 | 49 | **48** |
| LRM_Soybean | time | **0.018494** | 0.027114 | 0.027138 | 65.59715 | 53.06295 |
| | max | 7147213 | 275387 | 275376 | 250920 | **250908** |
| | min | 28248 | 240759 | 240799 | 250871 | 250859 |
| | max-min | 7118965 | 34628 | 34577 | **49** | **49** |
| LRM_Mouse | time | **0.022379** | 0.031497 | 0.032854 | 108.6254 | 76.35722 |
| | max | 30664387 | 779384 | 779437 | **580568** | 580585 |
| | min | 33565 | 527614 | 527628 | 580519 | 580537 |
| | max-min | 30630822 | 251770 | 251809 | 49 | **48** |
| LRM_hg38 | time | **0.020274** | 0.028600 | 0.029482 | 78.95541 | 61.48234 |
| | max | 55498448 | 961209 | 961201 | 558908 | **558895** |
| | min | 30505 | 466461 | 466485 | 558859 | 558846 |
| | max-min | 55467943 | 494748 | 494716 | **49** | **49** |
| RS_Pabies | time | **0.123976** | 0.148735 | 0.165093 | 1030.194 | 986.9083 |
| | max | 9648125 | 614677 | 614696 | **604182** | 604192 |
| | min | 102650 | 599520 | 599518 | 604132 | 604142 |
| | max-min | 9545475 | 15157 | 15178 | **50** | **50** |
| RS_Mouse | time | **0.006993** | 0.008741 | 0.008515 | 3.972343 | 3.719035 |
| | max | 1728720 | 141390 | 141585 | **129066** | 129095 |
| | min | 6133 | 121699 | 121709 | 129016 | 129048 |
| | max-min | 1722587 | 19691 | 19876 | 50 | **47** |
| RS_hg38 | time | **0.006199** | 0.007724 | 0.008358 | 3.341342 | 3.227035 |
| | max | 1852057 | 168929 | 169029 | **157252** | 157279 |
| | min | 5699 | 149202 | 149150 | 157201 | 157230 |
| | max-min | 1846358 | 19727 | 19879 | 51 | **49** |

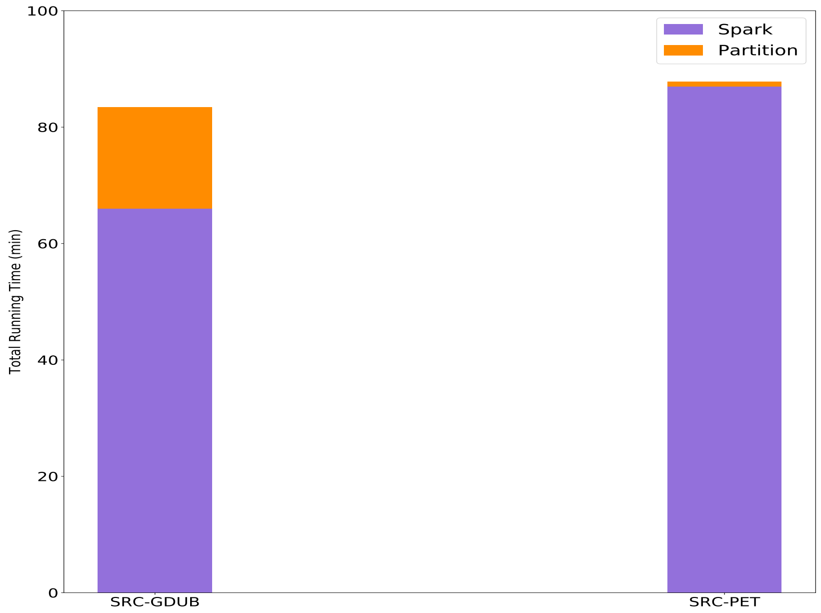


**Fig. S4.** The comparison of the running time of SRC-GDUB and SRC-PET based on RS_Pabies dataset. Note that Spark and Partition label represent the running time occupied by Spark and Partitioning during the whole process of SRC. SRC-GDUB and SRC-PET represent the SRC program using the GDUB and PET data partitioning method.

**3.3.4 CPU usage of RepeatClassifier** After the number of cores used in SRC is set to more than 60, the speedup ratio of threads decrease significantly. To figure out the reason, the CPU usage during the process of RepeatClassifier is summarized based on a small dataset. As shown in Fig. S8, RepeatClassifier runs in single-thread mode at first, and then uses 4 threads to run the program until the end. The maximum number of cores used will be close to the threshold of cores of Spark cluster when the number of cores in SRC is set to 60. This is the reason why the speedup ratio of SRC cannot reach the ′perfect′ line.

**3.3.5 Covering libraries of RepeatMasker** The Repeatmasker software (http://www.repeatmasker.org/), which makes use of *manually-curated* reference libraries of consensus sequences, has become the gold standard for masking. The libraries of RepeatMasker is consisted of RepBase and Dfam libraries. To prove that SRC has the same accuracy as that of RepeatClassifier, libraries from RepeatMasker are extracted using queryRepeatDatabase.pl. The extracted libraries are covered with the repetitive sequences classified by SRC and RepeatClassifier. The proportion and detailed classification of detection results of SRC and RepeatClas-

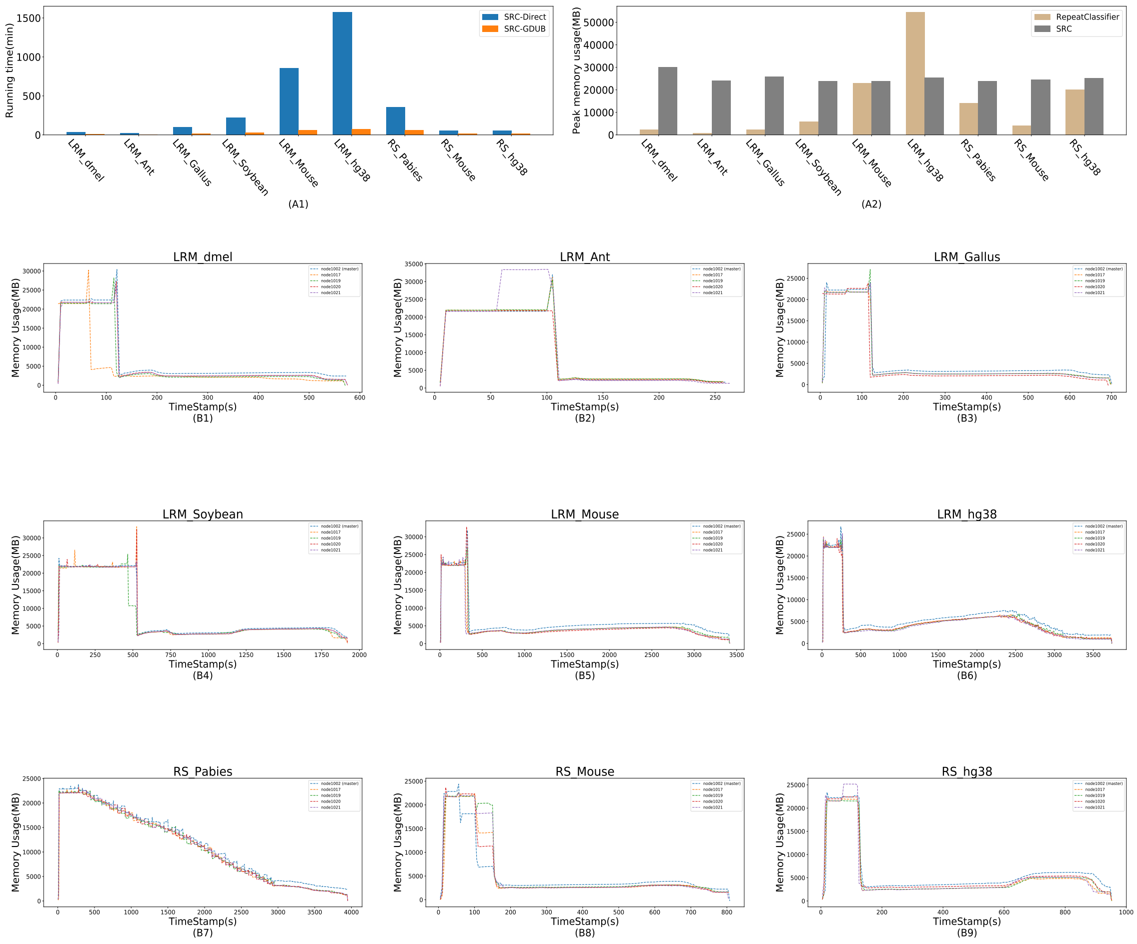


**Fig. S5.** The sub-graph(A1) shows the comparison of running time between SRC-GDUB and SRC-Direct. The sub-graph(A2) shows the peak memory usage of single node. The sub-graphs(B1-B9) show the memory usage of all nodes in the cluster.

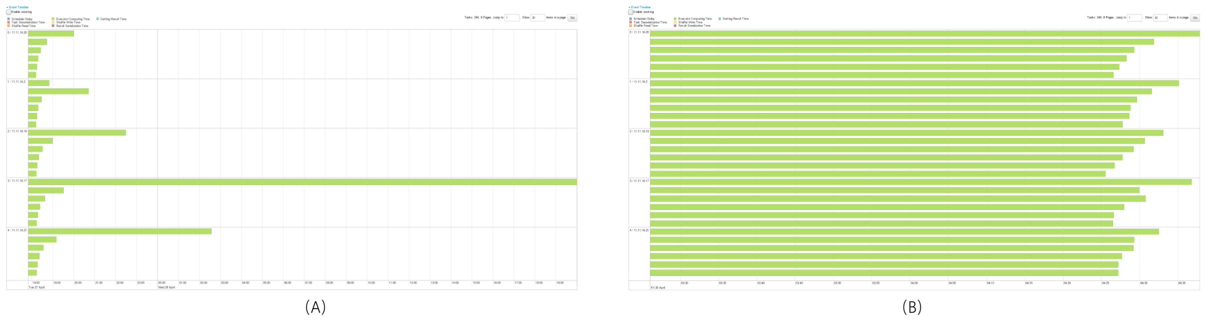


**Fig. S6.** Comparision of Spark event timelines between Direct and GDUB data partitioning methods based on Homo sapiens dataset. The sub-graph(A) shows the Spark event timeline using Direct data partitioning method. The sub-graph(B) shows the Spark event timeline using GDUB data partitioning method. Rectangles of green color in Sub-graph(A and B) represent the running time of each task in Spark.

sifier covering the libraries of RepBase and Dfam of all species datasets are shown in Table S4 to Table S12. It can be seen that SRC has the similar libraries coverage as RepeatClassifier.

**3.3.6 Covering reference genome** Repetitive sequences classified by SRC and RepeatClassifier are covered to the reference genome combining different types of alignment tools (Minimap2+BWA) which take the advantages of alignment in both short and long reads. The proportion and detailed classification of detection results of SRC and RepeatClassifier covering the repetitive regions on the reference genome of all species datasets are shown in Table S13 to S21. It can be seen that SRC has the similar reference coverage as RepeatClassifier.

**Table S4. The proportion and detailed classification of detection results of SRC and RepeatClassifier covering the libraries of RepBase and Dfam, which are generated by LongRepMarker using the reference genome of Drosophila melanogaster as input.**

| SparkRepeatClassifier | | | | RepeatClassifier | | |
|---|---|---|---|---|---|---|
| **sequence: 2489** | | | | **sequence: 2489** | | |
| **total length: 7220516bp** | | | | **total length: 7220516bp** | | |
| **bases masked: 3746452 bp ( 51.89%)** | | | | **bases masked: 3746647 bp ( 51.89%)** | | |
| **Repeat Types** | **Num of elements** | **Length occupied** | **Percentage of sequence** | **Num of elements** | **Length occupied** | **Percentage of sequence** |
| **DNA elements:** | 423 | 193711bp | 2.68% | 422 | 194158bp | 2.69% |
| –TcMar-Tigger: | 0 | 0bp | 0.00% | 0 | 0bp | 0.00% |
| –hAT-Charlie: | 0 | 0bp | 0.00% | 0 | 0bp | 0.00% |
| **LINEs:** | 1319 | 1043458bp | 14.45% | 1303 | 1032267bp | 14.30% |
| –L3/CR1: | 202 | 143230bp | 1.98% | 202 | 143230bp | 1.98% |
| –LINE1: | 0 | 0bp | 0.00% | 0 | 0bp | 0.00% |
| –LINE2: | 0 | 0bp | 0.00% | 0 | 0bp | 0.00% |
| **LTR elements:** | 2528 | 2356831bp | 32.64% | 2529 | 2357658bp | 32.65% |
| –ERVL: | 0 | 0bp | 0.00% | 0 | 0bp | 0.00% |
| –ERVL-MaLRs: | 0 | 0bp | 0.00% | 0 | 0bp | 0.00% |
| –ERV_classI: | 0 | 0bp | 0.00% | 0 | 0bp | 0.00% |
| –ERV_classII: | 0 | 0bp | 0.00% | 0 | 0bp | 0.00% |
| **Low complexity:** | 291 | 15714bp | 0.22% | 291 | 15714bp | 0.22% |
| **SINEs:** | 1 | 73bp | 0.00% | 2 | 208bp | 0.00% |
| –ALUs: | 0 | 0bp | 0.00% | 0 | 0bp | 0.00% |
| –MIRs: | 0 | 0bp | 0.00% | 0 | 0bp | 0.00% |
| **Satellites:** | 17 | 6719bp | 0.09% | 16 | 6584bp | 0.09% |
| **Simple repeats:** | 1108 | 74336bp | 1.03% | 1108 | 74336bp | 1.03% |
| **Small RNA:** | 29 | 13271bp | 0.18% | 29 | 13353bp | 0.18% |
| **Total interspersed repeats:** | | 3646136bp | 50.50% | | 3648025bp | 50.52% |
| **Unclassified:** | 142 | 52063bp | 0.72% | 164 | 63734bp | 0.88% |

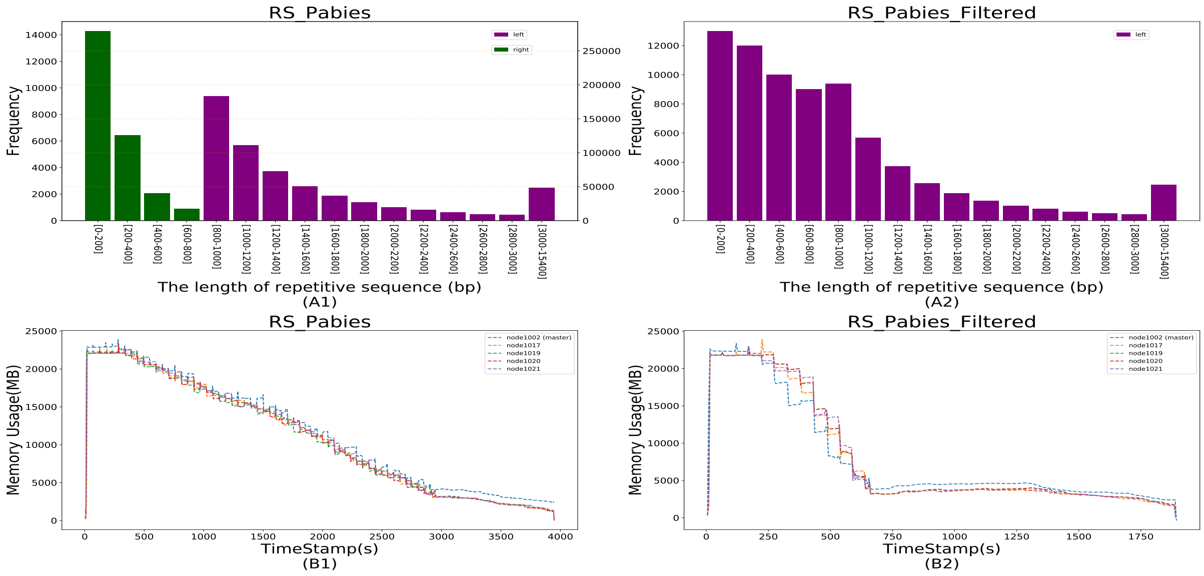


**Fig. S7.** The relationship between memory usage and sequences length distribution. The sub-graphs(A1-A2) show the length distribution of repetitive sequences. The sub-graphs(B1-B2) show the memory usage of all nodes in the cluster. Note that two y-axes exist in sub-graphs(A1) since the frequency difference of the dataset is too large to be represented by only one y-axis. Purple bars correspond to the left y axis, which mean the frequency of repetitive sequence <= 10000, and green bars correspond to the right y axis, which mean the frequency of repetitive sequence > 10000.

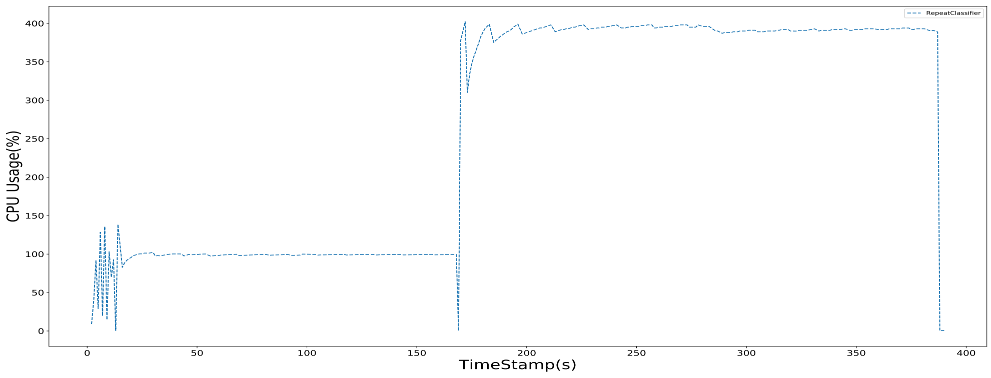


**Fig. S8.** CPU usage of RepeatClassifier. Note that 400 on the y-axis means that the program uses 4 threads.

**Table S5. The proportion and detailed classification of detection results of SRC and RepeatClassifier covering the libraries of RepBase and Dfam, which are generated by LongRepMarker using the reference genome of Acromyrmex echinatior as input.**

| SparkRepeatClassifier | | | | RepeatClassifier | | |
|---|---|---|---|---|---|---|
| **sequence: 254** | | | | **sequence: 254** | | |
| **total length: 214457bp** | | | | **total length: 214457bp** | | |
| **bases masked: 168914 bp ( 78.76%)** | | | | **bases masked: 168914 bp ( 78.76%)** | | |
| **Repeat Types** | **Num of elements** | **Length occupied** | **Percentage of sequence** | **Num of elements** | **Length occupied** | **Percentage of sequence** |
| **DNA elements:** | 97 | 75203bp | 35.07% | 98 | 75226bp | 35.08% |
| –TcMar-Tigger: | 0 | 0bp | 0.00% | 0 | 0bp | 0.00% |
| –hAT-Charlie: | 0 | 0bp | 0.00% | 0 | 0bp | 0.00% |
| **LINEs:** | 19 | 8528bp | 3.98% | 19 | 8528bp | 3.98% |
| –L3/CR1: | 0 | 0bp | 0.00% | 0 | 0bp | 0.00% |
| –LINE1: | 0 | 0bp | 0.00% | 0 | 0bp | 0.00% |
| –LINE2: | 0 | 0bp | 0.00% | 0 | 0bp | 0.00% |
| **LTR elements:** | 71 | 48344bp | 22.54% | 75 | 48344bp | 22.54% |
| –ERVL: | 0 | 0bp | 0.00% | 0 | 0bp | 0.00% |
| –ERVL-MaLRs: | 0 | 0bp | 0.00% | 0 | 0bp | 0.00% |
| –ERV_classI: | 0 | 0bp | 0.00% | 0 | 0bp | 0.00% |
| –ERV_classII: | 0 | 0bp | 0.00% | 0 | 0bp | 0.00% |
| **Low complexity:** | 2 | 110bp | 0.05% | 2 | 110bp | 0.05% |
| **SINEs:** | 1 | 69bp | 0.03% | 1 | 69bp | 0.03% |
| –ALUs: | 0 | 0bp | 0.00% | 0 | 0bp | 0.00% |
| –MIRs: | 0 | 0bp | 0.00% | 0 | 0bp | 0.00% |
| **Satellites:** | 0 | 0bp | 0.00% | 0 | 0bp | 0.00% |
| **Simple repeats:** | 185 | 30494bp | 14.22% | 185 | 30494bp | 14.22% |
| **Small RNA:** | 6 | 566bp | 0.26% | 6 | 566bp | 0.26% |
| **Total interspersed repeats:** | | 138083bp | 64.39% | | 138106bp | 64.40% |
| **Unclassified:** | 33 | 5939bp | 2.77% | 31 | 5939bp | 2.77% |

**Table S6. The proportion and detailed classification of detection results of SRC and RepeatClassifier covering the libraries of RepBase and Dfam, which are generated by LongRepMarker using the reference genome of Gallus gallusbankiva as input.**

| SparkRepeatClassifier | | | | RepeatClassifier | | |
|---|---|---|---|---|---|---|
| **sequence: 512** | | | | **sequence: 512** | | |
| **total length: 362626bp** | | | | **total length: 362626bp** | | |
| **bases masked: 255267 bp ( 70.39%)** | | | | **bases masked: 255267 bp ( 70.39%)** | | |
| **Repeat Types** | **Num of elements** | **Length occupied** | **Percentage of sequence** | **Num of elements** | **Length occupied** | **Percentage of sequence** |
| **DNA elements:** | 18 | 3974bp | 1.10% | 18 | 3974bp | 1.10% |
| –TcMar-Tigger: | 0 | 0bp | 0.00% | 0 | 0bp | 0.00% |
| –hAT-Charlie: | 2 | 346bp | 0.10% | 2 | 346bp | 0.10% |
| **LINEs:** | 163 | 119316bp | 32.90% | 162 | 119130bp | 32.85% |
| –L3/CR1: | 163 | 119316bp | 32.90% | 162 | 119130bp | 32.85% |
| –LINE1: | 0 | 0bp | 0.00% | 0 | 0bp | 0.00% |
| –LINE2: | 0 | 0bp | 0.00% | 0 | 0bp | 0.00% |
| **LTR elements:** | 70 | 79597bp | 21.95% | 70 | 79783bp | 22.00% |
| –ERVL: | 40 | 45157bp | 12.45% | 39 | 44738bp | 12.34% |
| –ERVL-MaLRs: | 0 | 0bp | 0.00% | 0 | 0bp | 0.00% |
| –ERV_classI: | 7 | 9997bp | 2.76% | 7 | 9997bp | 2.76% |
| –ERV_classII: | 22 | 23497bp | 6.48% | 23 | 24102bp | 6.65% |
| **Low complexity:** | 3 | 147bp | 0.04% | 3 | 147bp | 0.04% |
| **SINEs:** | 9 | 784bp | 0.22% | 9 | 784bp | 0.22% |
| –ALUs: | 0 | 0bp | 0.00% | 0 | 0bp | 0.00% |
| –MIRs: | 0 | 0bp | 0.00% | 0 | 0bp | 0.00% |
| **Satellites:** | 5 | 5709bp | 1.57% | 5 | 5709bp | 1.57% |
| **Simple repeats:** | 197 | 31810bp | 8.77% | 197 | 31810bp | 8.77% |
| **Small RNA:** | 39 | 6135bp | 1.69% | 39 | 6135bp | 1.69% |
| **Total interspersed repeats:** | | 212012bp | 58.47% | | 212012bp | 58.47% |
| **Unclassified:** | 6 | 8341bp | 2.30% | 6 | 8341bp | 2.30% |

**Table S7. The proportion and detailed classification of detection results of SRC and RepeatClassifier covering the libraries of RepBase and Dfam, which are generated by LongRepMarker using the reference genome of Glycine max as input.**

| SparkRepeatClassifier | | | | RepeatClassifier | | |
|---|---|---|---|---|---|---|
| **sequence: 758** | | | | **sequence: 758** | | |
| **total length: 1646292bp** | | | | **total length: 1646292bp** | | |
| **bases masked: 1536173 bp ( 93.31%)** | | | | **bases masked: 1536173 bp ( 93.31%)** | | |
| **Repeat Types** | **Num of elements** | **Length occupied** | **Percentage of sequence** | **Num of elements** | **Length occupied** | **Percentage of sequence** |
| **DNA elements:** | 131 | 154230bp | 9.37% | 131 | 154230bp | 9.37% |
| –TcMar-Tigger: | 0 | 0bp | 0.00% | 0 | 0bp | 0.00% |
| –hAT-Charlie: | 0 | 0bp | 0.00% | 0 | 0bp | 0.00% |
| **LINEs:** | 45 | 80754bp | 4.91% | 45 | 80754bp | 4.91% |
| –L3/CR1: | 0 | 0bp | 0.00% | 0 | 0bp | 0.00% |
| –LINE1: | 44 | 77578bp | 4.71% | 44 | 77578bp | 4.71% |
| –LINE2: | 0 | 0bp | 0.00% | 0 | 0bp | 0.00% |
| **LTR elements:** | 881 | 1238562bp | 75.23% | 884 | 1238476bp | 75.23% |
| –ERVL: | 0 | 0bp | 0.00% | 0 | 0bp | 0.00% |
| –ERVL-MaLRs: | 0 | 0bp | 0.00% | 0 | 0bp | 0.00% |
| –ERV_classI: | 0 | 0bp | 0.00% | 0 | 0bp | 0.00% |
| –ERV_classII: | 0 | 0bp | 0.00% | 0 | 0bp | 0.00% |
| **Low complexity:** | 9 | 1018bp | 0.06% | 9 | 1018bp | 0.06% |
| **SINEs:** | 1 | 18bp | 0.00% | 1 | 18bp | 0.00% |
| –ALUs: | 0 | 0bp | 0.00% | 0 | 0bp | 0.00% |
| –MIRs: | 0 | 0bp | 0.00% | 0 | 0bp | 0.00% |
| **Satellites:** | 0 | 0bp | 0.00% | 2 | 86bp | 0.01% |
| **Simple repeats:** | 200 | 31493bp | 1.91% | 200 | 31493bp | 1.91% |
| **Small RNA:** | 22 | 6625bp | 0.40% | 22 | 6625bp | 0.40% |
| **Total interspersed repeats:** | | 1498583bp | 91.03% | | 1498497bp | 91.02% |
| **Unclassified:** | 27 | 25019bp | 1.52% | 27 | 25019bp | 1.52% |

**Table S8. The proportion and detailed classification of detection results of SRC and RepeatClassifier covering the libraries of RepBase and Dfam, which are generated by LongRepMarker using the reference genome of Mus musculus as input.**

| SparkRepeatClassifier | | | | RepeatClassifier | | |
|---|---|---|---|---|---|---|
| **sequence: 1561** | | | | **sequence: 1561** | | |
| **total length: 1680566bp** | | | | **total length: 1680566bp** | | |
| **bases masked: 1044497 bp ( 62.15%)** | | | | **bases masked: 1044654 bp ( 62.16%)** | | |
| **Repeat Types** | **Num of elements** | **Length occupied** | **Percentage of sequence** | **Num of elements** | **Length occupied** | **Percentage of sequence** |
| **DNA elements:** | 57 | 7136bp | 0.42% | 57 | 6916bp | 0.41% |
| –TcMar-Tigger: | 9 | 1410bp | 0.08% | 9 | 1410bp | 0.08% |
| –hAT-Charlie: | 32 | 3880bp | 0.23% | 32 | 3660bp | 0.22% |
| **LINEs:** | 825 | 585528bp | 34.84% | 809 | 582427bp | 34.66% |
| –L3/CR1: | 0 | 0bp | 0.00% | 0 | 0bp | 0.00% |
| –LINE1: | 823 | 583297bp | 34.71% | 807 | 580196bp | 34.52% |
| –LINE2: | 1 | 94bp | 0.01% | 1 | 94bp | 0.01% |
| **LTR elements:** | 493 | 346837bp | 20.64% | 500 | 349619bp | 20.80% |
| –ERVL: | 85 | 56340bp | 3.35% | 86 | 56485bp | 3.36% |
| –ERVL-MaLRs: | 57 | 10909bp | 0.65% | 56 | 10906bp | 0.65% |
| –ERV_classI: | 78 | 67482bp | 4.02% | 77 | 67566bp | 4.02% |
| –ERV_classII: | 265 | 209734bp | 12.48% | 270 | 211757bp | 12.60% |
| **Low complexity:** | 35 | 1618bp | 0.10% | 35 | 1618bp | 0.10% |
| **SINEs:** | 325 | 62379bp | 3.71% | 325 | 62478bp | 3.72% |
| –ALUs: | 272 | 52730bp | 3.14% | 273 | 52891bp | 3.15% |
| –MIRs: | 8 | 1464bp | 0.09% | 8 | 1464bp | 0.09% |
| **Satellites:** | 8 | 4208bp | 0.25% | 8 | 4208bp | 0.25% |
| **Simple repeats:** | 314 | 36351bp | 2.16% | 314 | 36351bp | 2.16% |
| **Small RNA:** | 29 | 3693bp | 0.22% | 32 | 3897bp | 0.23% |
| **Total interspersed repeats:** | | 1014113bp | 60.34% | | 1013374bp | 60.30% |
| **Unclassified:** | 40 | 12233bp | 0.73% | 38 | 11934bp | 0.71% |

**Table S9. The proportion and detailed classification of detection results of SRC and RepeatClassifier covering the libraries of RepBase and Dfam, which are generated by LongRepMarker using the reference genome of Homo sapiens as input.**

| SparkRepeatClassifier | | | | RepeatClassifier | | |
|---|---|---|---|---|---|---|
| **sequence: 1512** | | | | **sequence: 1512** | | |
| **total length: 1647075bp** | | | | **total length: 1647075bp** | | |
| **bases masked: 1358054 bp ( 82.45%)** | | | | **bases masked: 1358050 bp ( 82.45%)** | | |
| **Repeat Types** | **Num of elements** | **Length occupied** | **Percentage of sequence** | **Num of elements** | **Length occupied** | **Percentage of sequence** |
| **DNA elements:** | 110 | 25838bp | 1.57% | 111 | 25814bp | 1.57% |
| –TcMar-Tigger: | 35 | 11048bp | 0.67% | 37 | 11048bp | 0.67% |
| –hAT-Charlie: | 41 | 8781bp | 0.53% | 40 | 8757bp | 0.53% |
| **LINEs:** | 1376 | 690975bp | 41.95% | 1332 | 676584bp | 41.08% |
| –L3/CR1: | 5 | 708bp | 0.04% | 5 | 723bp | 0.04% |
| –LINE1: | 1337 | 682454bp | 41.43% | 1297 | 668428bp | 40.58% |
| –LINE2: | 11 | 1455bp | 0.09% | 13 | 1570bp | 0.10% |
| **LTR elements:** | 566 | 327086bp | 19.86% | 571 | 328870bp | 19.97% |
| –ERVL: | 98 | 39268bp | 2.38% | 102 | 40575bp | 2.46% |
| –ERVL-MaLRs: | 32 | 8118bp | 0.49% | 36 | 8525bp | 0.52% |
| –ERV_classI: | 370 | 220447bp | 13.38% | 368 | 220879bp | 13.41% |
| –ERV_classII: | 54 | 57466bp | 3.49% | 53 | 57104bp | 3.47% |
| **Low complexity:** | 11 | 483bp | 0.03% | 11 | 483bp | 0.03% |
| **SINEs:** | 709 | 255186bp | 15.49% | 740 | 262816bp | 15.96% |
| –ALUs: | 690 | 251356bp | 15.26% | 720 | 258865bp | 15.72% |
| –MIRs: | 17 | 3552bp | 0.22% | 17 | 3552bp | 0.22% |
| **Satellites:** | 24 | 10205bp | 0.62% | 24 | 10205bp | 0.62% |
| **Simple repeats:** | 216 | 31821bp | 1.93% | 216 | 31821bp | 1.93% |
| **Small RNA:** | 14 | 1276bp | 0.08% | 19 | 2841bp | 0.17% |
| **Total interspersed repeats:** | | 1356298bp | 82.35% | | 1354172bp | 82.22% |
| **Unclassified:** | 185 | 57213bp | 3.47% | 200 | 60088bp | 3.65% |

**Table S10. The proportion and detailed classification of detection results of SRC and RepeatClassifier covering the libraries of RepBase and Dfam, which are generated by RepeatScout using the reference genome of Picea abies as input.**

| SparkRepeatClassifier | | | | RepeatClassifier | | |
|---|---|---|---|---|---|---|
| **sequence: 445** | | | | **sequence: 445** | | |
| **total length: 1039165bp** | | | | **total length: 1039165bp** | | |
| **bases masked: 992982 bp ( 95.56%)** | | | | **bases masked: 992982 bp ( 95.56%)** | | |
| **Repeat Types** | **Num of elements** | **Length occupied** | **Percentage of sequence** | **Num of elements** | **Length occupied** | **Percentage of sequence** |
| **DNA elements:** | 0 | 0bp | 0.00% | 0 | 0bp | 0.00% |
| –TcMar-Tigger: | 0 | 0bp | 0.00% | 0 | 0bp | 0.00% |
| –hAT-Charlie: | 0 | 0bp | 0.00% | 0 | 0bp | 0.00% |
| **LINEs:** | 9 | 24333bp | 2.34% | 9 | 24333bp | 2.34% |
| –L3/CR1: | 0 | 0bp | 0.00% | 0 | 0bp | 0.00% |
| –LINE1: | 9 | 24333bp | 2.34% | 9 | 24333bp | 2.34% |
| –LINE2: | 0 | 0bp | 0.00% | 0 | 0bp | 0.00% |
| **LTR elements:** | 409 | 934061bp | 89.89% | 401 | 934061bp | 89.89% |
| –ERVL: | 0 | 0bp | 0.00% | 0 | 0bp | 0.00% |
| –ERVL-MaLRs: | 0 | 0bp | 0.00% | 0 | 0bp | 0.00% |
| –ERV_classI: | 0 | 0bp | 0.00% | 0 | 0bp | 0.00% |
| –ERV_classII: | 0 | 0bp | 0.00% | 0 | 0bp | 0.00% |
| **Low complexity:** | 0 | 0bp | 0.00% | 0 | 0bp | 0.00% |
| **SINEs:** | 0 | 0bp | 0.00% | 0 | 0bp | 0.00% |
| –ALUs: | 0 | 0bp | 0.00% | 0 | 0bp | 0.00% |
| –MIRs: | 0 | 0bp | 0.00% | 0 | 0bp | 0.00% |
| **Satellites:** | 0 | 0bp | 0.00% | 0 | 0bp | 0.00% |
| **Simple repeats:** | 178 | 30036bp | 2.89% | 178 | 30036bp | 2.89% |
| **Small RNA:** | 18 | 4504bp | 0.43% | 18 | 4504bp | 0.43% |
| **Total interspersed repeats:** | | 958442bp | 92.23% | | 958442bp | 92.23% |
| **Unclassified:** | 1 | 48bp | 0.00% | 1 | 48bp | 0.00% |

**Table S11. The proportion and detailed classification of detection results of SRC and RepeatClassifier covering the libraries of RepBase and Dfam, which are generated by RepeatScout using the reference genome of Mus musculus as input.**

| SparkRepeatClassifier | | | | RepeatClassifier | | |
|---|---|---|---|---|---|---|
| **sequence: 1561** | | | | **sequence: 1561** | | |
| **total length: 1680566bp** | | | | **total length: 1680566bp** | | |
| **bases masked: 989911 bp ( 58.90%)** | | | | **bases masked: 989953 bp ( 58.91%)** | | |
| **Repeat Types** | **Num of elements** | **Length occupied** | **Percentage of sequence** | **Num of elements** | **Length occupied** | **Percentage of sequence** |
| **DNA elements:** | 40 | 5486bp | 0.33% | 33 | 4009bp | 0.24% |
| –TcMar-Tigger: | 1 | 107bp | 0.01% | 1 | 107bp | 0.01% |
| –hAT-Charlie: | 31 | 4364bp | 0.26% | 25 | 3117bp | 0.19% |
| **LINEs:** | 462 | 415254bp | 24.71% | 476 | 418545bp | 24.91% |
| –L3/CR1: | 0 | 0bp | 0.00% | 0 | 0bp | 0.00% |
| –LINE1: | 458 | 415049bp | 24.70% | 471 | 417701bp | 24.85% |
| –LINE2: | 4 | 205bp | 0.01% | 4 | 205bp | 0.01% |
| **LTR elements:** | 663 | 464631bp | 27.65% | 660 | 459383bp | 27.34% |
| –ERVL: | 84 | 55567bp | 3.31% | 82 | 54287bp | 3.23% |
| –ERVL-MaLRs: | 70 | 17508bp | 1.04% | 71 | 16714bp | 0.99% |
| –ERV_classI: | 127 | 90204bp | 5.37% | 118 | 87098bp | 5.18% |
| –ERV_classII: | 381 | 293900bp | 17.49% | 384 | 292250bp | 17.39% |
| **Low complexity:** | 40 | 1873bp | 0.11% | 40 | 1873bp | 0.11% |
| **SINEs:** | 270 | 55810bp | 3.32% | 287 | 60244bp | 3.58% |
| –ALUs: | 200 | 45464bp | 2.71% | 214 | 50159bp | 2.98% |
| –MIRs: | 5 | 423bp | 0.03% | 5 | 439bp | 0.03% |
| **Satellites:** | 14 | 5702bp | 0.34% | 14 | 5063bp | 0.30% |
| **Simple repeats:** | 327 | 36959bp | 2.20% | 327 | 36959bp | 2.20% |
| **Small RNA:** | 55 | 4831bp | 0.29% | 55 | 4831bp | 0.29% |
| **Total interspersed repeats:** | | 951264bp | 56.60% | | 951831bp | 56.64% |
| **Unclassified:** | 37 | 10083bp | 0.60% | 34 | 9650bp | 0.57% |

**Table S12. The proportion and detailed classification of detection results of SRC and RepeatClassifier covering the libraries of RepBase and Dfam, which are generated by RepeatScout using the reference genome of Homo sapiens as input.**

| SparkRepeatClassifier | | | | RepeatClassifier | | |
|---|---|---|---|---|---|---|
| **sequence: 1512** | | | | **sequence: 1512** | | |
| **total length: 1647075bp** | | | | **total length: 1647075bp** | | |
| **bases masked: 1215893 bp ( 73.82%)** | | | | **bases masked: 1215905 bp ( 73.82%)** | | |
| **Repeat Types** | **Num of elements** | **Length occupied** | **Percentage of sequence** | **Num of elements** | **Length occupied** | **Percentage of sequence** |
| **DNA elements:** | 220 | 63559bp | 3.86% | 224 | 64597bp | 3.92% |
| –TcMar-Tigger: | 76 | 32881bp | 2.00% | 86 | 34940bp | 2.12% |
| –hAT-Charlie: | 48 | 10950bp | 0.66% | 48 | 11006bp | 0.67% |
| **LINEs:** | 582 | 293763bp | 17.84% | 618 | 297993bp | 18.09% |
| –L3/CR1: | 5 | 818bp | 0.05% | 4 | 805bp | 0.05% |
| –LINE1: | 564 | 289618bp | 17.58% | 602 | 294357bp | 17.87% |
| –LINE2: | 10 | 3013bp | 0.18% | 9 | 2517bp | 0.15% |
| **LTR elements:** | 920 | 574392bp | 34.87% | 907 | 572581bp | 34.76% |
| –ERVL: | 158 | 88400bp | 5.37% | 152 | 87221bp | 5.30% |
| –ERVL-MaLRs: | 41 | 12780bp | 0.78% | 47 | 13970bp | 0.85% |
| –ERV_classI: | 647 | 392001bp | 23.80% | 638 | 389747bp | 23.66% |
| –ERV_classII: | 60 | 78798bp | 4.78% | 56 | 79131bp | 4.80% |
| **Low complexity:** | 17 | 779bp | 0.05% | 17 | 779bp | 0.05% |
| **SINEs:** | 695 | 180875bp | 10.98% | 684 | 188307bp | 11.43% |
| –ALUs: | 677 | 179017bp | 10.87% | 670 | 187083bp | 11.36% |
| –MIRs: | 12 | 1060bp | 0.06% | 9 | 613bp | 0.04% |
| **Satellites:** | 38 | 14435bp | 0.88% | 38 | 14435bp | 0.88% |
| **Simple repeats:** | 257 | 34176bp | 2.07% | 257 | 34176bp | 2.07% |
| **Small RNA:** | 54 | 12449bp | 0.76% | 54 | 12449bp | 0.76% |
| **Total interspersed repeats:** | | 1172834bp | 71.21% | | 1178001bp | 71.52% |
| **Unclassified:** | 161 | 60245bp | 3.66% | 142 | 54523bp | 3.31% |

**Table S13. The proportion and detailed classification of detection results of SRC and RepeatClassifier covering the repetitive regions, which are generated by LongRepMarker using the reference genome of Drosophila melanogaster as input.**

| SparkRepeatClassifier | | | | RepeatClassifier | | |
|---|---|---|---|---|---|---|
| **sequence: 15** | | | | **sequence: 15** | | |
| **total length: 168736537bp** | | | | **total length: 168736537bp** | | |
| **bases masked: 34098031 bp (20.21%)** | | | | **bases masked: 34097818 bp (20.21%)** | | |
| **Repeat Types** | **Num of elements** | **Length occupied** | **Percentage of sequence** | **Num of elements** | **Length occupied** | **Percentage of sequence** |
| **DNA:** | 14298 | 1746619bp | 1.04% | 14300 | 1746454bp | 1.04% |
| –Academ-1: | 6 | 3667bp | 0.00% | 6 | 3667bp | 0.00% |
| –CMC-EnSpm: | 9 | 3415bp | 0.00% | 9 | 3415bp | 0.00% |
| –CMC-Transib: | 1448 | 153013bp | 0.09% | 1450 | 153013bp | 0.09% |
| –MULE-MuDR: | 12 | 7323bp | 0.00% | 12 | 7323bp | 0.00% |
| –MULE-NOF: | 726 | 100063bp | 0.06% | 726 | 100063bp | 0.06% |
| –Maverick: | 14 | 6413bp | 0.00% | 14 | 6413bp | 0.00% |
| –MuLE-NOF: | 6 | 239bp | 0.00% | 6 | 239bp | 0.00% |
| –OTHER: | 206 | 22100bp | 0.01% | 206 | 22100bp | 0.01% |
| –P: | 6582 | 935918bp | 0.55% | 6582 | 935918bp | 0.55% |
| –PIF-Harbinger: | 16 | 894bp | 0.00% | 16 | 894bp | 0.00% |
| –PiggyBac: | 19 | 955bp | 0.00% | 19 | 955bp | 0.00% |
| –TcMar-Pogo: | 717 | 54830bp | 0.03% | 721 | 54830bp | 0.03% |
| –TcMar-Tc1: | 2423 | 260265bp | 0.15% | 2423 | 260265bp | 0.15% |
| –hAT-Ac: | 823 | 82579bp | 0.05% | 823 | 82579bp | 0.05% |
| –hAT-Tag1: | 61 | 14006bp | 0.01% | 61 | 14006bp | 0.01% |
| –hAT-Tip100: | 25 | 14147bp | 0.01% | 25 | 14147bp | 0.01% |
| –hAT-hATm: | 13 | 9519bp | 0.01% | 13 | 9519bp | 0.01% |
| –hAT-hobo: | 1192 | 133839bp | 0.08% | 1188 | 133674bp | 0.08% |
| **LINE:** | 63010 | 7968013bp | 4.72% | 62882 | 7985953bp | 4.73% |
| –CR1: | 3231 | 604771bp | 0.36% | 3235 | 605943bp | 0.36% |
| –I: | 1627 | 302608bp | 0.18% | 1627 | 302608bp | 0.18% |
| –I-Jockey: | 17000 | 2648503bp | 1.57% | 16848 | 2665316bp | 1.58% |
| –Jockey: | 16801 | 2866297bp | 1.70% | 16908 | 2821244bp | 1.67% |
| –LOA: | 1108 | 166353bp | 0.10% | 1026 | 167430bp | 0.10% |
| –OTHER: | 127 | 94706bp | 0.06% | 127 | 94706bp | 0.06% |
| –R1: | 19563 | 2618643bp | 1.55% | 19563 | 2618643bp | 1.55% |
| –R1-LOA: | 448 | 130728bp | 0.08% | 443 | 128628bp | 0.08% |
| –R2: | 3087 | 205770bp | 0.12% | 3087 | 205770bp | 0.12% |
| –R2-NeSL: | 8 | 617bp | 0.00% | 8 | 617bp | 0.00% |
| –RTE-X: | 10 | 17039bp | 0.01% | 10 | 17039bp | 0.01% |
| **LTR:** | 127541 | 15911903bp | 9.43% | 127528 | 15912624bp | 9.43% |
| –Copia: | 7021 | 961221bp | 0.57% | 7017 | 960711bp | 0.57% |
| –ERVK: | 15 | 6458bp | 0.00% | 15 | 6458bp | 0.00% |
| –Gypsy: | 83678 | 11517560bp | 6.83% | 83669 | 11517206bp | 6.83% |
| –Gypsy-Cigr: | 5 | 559bp | 0.00% | 5 | 559bp | 0.00% |
| –OTHER: | 83 | 70018bp | 0.04% | 83 | 70018bp | 0.04% |
| –Pao: | 35945 | 3426976bp | 2.03% | 35945 | 3428561bp | 2.03% |
| –Viper: | 794 | 34755bp | 0.02% | 794 | 34755bp | 0.02% |
| **Other:** | 7666 | 660761bp | 0.39% | 7666 | 660761bp | 0.39% |
| –OTHER: | 7666 | 660761bp | 0.39% | 7666 | 660761bp | 0.39% |
| **RC:** | 1457 | 306200bp | 0.18% | 1457 | 306200bp | 0.18% |
| –Helitron: | 1457 | 306200bp | 0.18% | 1457 | 306200bp | 0.18% |
| **RNA:** | 116 | 20947bp | 0.01% | 116 | 20947bp | 0.01% |
| –OTHER: | 116 | 20947bp | 0.01% | 116 | 20947bp | 0.01% |
| **SINE:** | 236 | 31505bp | 0.02% | 236 | 31505bp | 0.02% |
| –5S: | 98 | 25877bp | 0.02% | 98 | 25877bp | 0.02% |
| –ID: | 35 | 929bp | 0.00% | 35 | 929bp | 0.00% |
| –U: | 31 | 2135bp | 0.00% | 31 | 2135bp | 0.00% |
| –tRNA: | 31 | 1468bp | 0.00% | 31 | 1468bp | 0.00% |
| –tRNA-RTE: | 41 | 1096bp | 0.00% | 41 | 1096bp | 0.00% |
| **Satellite:** | 25288 | 3682739bp | 2.18% | 25288 | 3682739bp | 2.18% |
| –OTHER: | 25288 | 3682739bp | 2.18% | 25288 | 3682739bp | 2.18% |
| **Simple:** | 4161 | 194894bp | 0.12% | 4161 | 194894bp | 0.12% |
| –repeat: | 4161 | 194894bp | 0.12% | 4161 | 194894bp | 0.12% |
| **Unknown:** | 42126 | 4185948bp | 2.48% | 42257 | 4231252bp | 2.51% |
| –OTHER: | 42126 | 4185948bp | 2.48% | 42257 | 4231252bp | 2.51% |
| **rRNA:** | 32958 | 1783949bp | 1.06% | 32958 | 1783949bp | 1.06% |
| –OTHER: | 32958 | 1783949bp | 1.06% | 32958 | 1783949bp | 1.06% |
| **snRNA:** | 34 | 1869bp | 0.00% | 34 | 1869bp | 0.00% |
| –OTHER: | 34 | 1869bp | 0.00% | 34 | 1869bp | 0.00% |
| **tRNA:** | 937 | 23307bp | 0.01% | 937 | 23307bp | 0.01% |
| –OTHER: | 937 | 23307bp | 0.01% | 937 | 23307bp | 0.01% |

**Table S14. The proportion and detailed classification of detection results of SRC and RepeatClassifier covering the repetitive regions, which are generated by LongRepMarker using the reference genome of Acromyrmex echinatior as input.**

| | SparkRepeatClassifier | | | RepeatClassifier | | |
|---|---|---|---|---|---|---|
| **sequence: 4339** | | | | **sequence: 4339** | | |
| **total length: 295944863bp** | | | | **total length: 295944863bp** | | |
| **bases masked: 11663344 bp (3.94%)** | | | | **bases masked: 11662860 bp (3.94%)** | | |
| **Repeat Types** | **Num of elements** | **Length occupied** | **Percentage of sequence** | **Num of elements** | **Length occupied** | **Percentage of sequence** |
| **DNA:** | 61552 | 4928388bp | 1.67% | 61498 | 4927684bp | 1.67% |
| –Academ-1: | 5 | 661bp | 0.00% | 5 | 661bp | 0.00% |
| –CMC-Chapaev-3: | 188 | 28829bp | 0.01% | 188 | 28829bp | 0.01% |
| –CMC-EnSpm: | 150 | 44955bp | 0.02% | 150 | 44955bp | 0.02% |
| –CMC-Transib: | 13 | 862bp | 0.00% | 13 | 862bp | 0.00% |
| –Crypton-V: | 40 | 4320bp | 0.00% | 40 | 4320bp | 0.00% |
| –IS3EU: | 8 | 3241bp | 0.00% | 8 | 3241bp | 0.00% |
| –Kolobok-Hydra: | 904 | 76209bp | 0.03% | 897 | 75783bp | 0.03% |
| –Kolobok-T2: | 8759 | 585078bp | 0.20% | 8759 | 585078bp | 0.20% |
| –MULE-NOF: | 18 | 5612bp | 0.00% | 18 | 5612bp | 0.00% |
| –Maverick: | 15263 | 860598bp | 0.29% | 15222 | 860259bp | 0.29% |
| –Merlin: | 69 | 21185bp | 0.01% | 69 | 21185bp | 0.01% |
| –MuLE-NOF: | 4 | 2252bp | 0.00% | 4 | 2252bp | 0.00% |
| –MuLE-NOF?: | 6 | 593bp | 0.00% | 6 | 593bp | 0.00% |
| –OTHER: | 661 | 179910bp | 0.06% | 669 | 181756bp | 0.06% |
| –P: | 34 | 2854bp | 0.00% | 34 | 2854bp | 0.00% |
| –PIF-Harbinger: | 15 | 1743bp | 0.00% | 15 | 1743bp | 0.00% |
| –PIF-Spy: | 18 | 1991bp | 0.00% | 18 | 1991bp | 0.00% |
| –PiggyBac: | 63 | 8101bp | 0.00% | 63 | 8101bp | 0.00% |
| –TcMar: | 343 | 28338bp | 0.01% | 343 | 28338bp | 0.01% |
| –TcMar-Cweed: | 31 | 4601bp | 0.00% | 31 | 4601bp | 0.00% |
| –TcMar-Fot1: | 24 | 14562bp | 0.00% | 24 | 14562bp | 0.00% |
| –TcMar-Mariner: | 22053 | 2085501bp | 0.70% | 22041 | 2084881bp | 0.70% |
| –TcMar-Tc1: | 11262 | 892895bp | 0.30% | 11261 | 892895bp | 0.30% |
| –TcMar-Tc4: | 1184 | 63484bp | 0.02% | 1184 | 63484bp | 0.02% |
| –TcMar-Tigger: | 5 | 434bp | 0.00% | 5 | 434bp | 0.00% |
| –hAT: | 79 | 9856bp | 0.00% | 79 | 9856bp | 0.00% |
| –hAT-Ac: | 47 | 19373bp | 0.01% | 47 | 19373bp | 0.01% |
| –hAT-Blackjack: | 210 | 19502bp | 0.01% | 210 | 19502bp | 0.01% |
| –hAT-Charlie: | 70 | 28741bp | 0.01% | 69 | 28569bp | 0.01% |
| –hAT-Pegasus: | 12 | 7622bp | 0.00% | 12 | 7622bp | 0.00% |
| –hAT-Tip100: | 14 | 6534bp | 0.00% | 14 | 6534bp | 0.00% |
| **LINE:** | 9011 | 812146bp | 0.27% | 9005 | 812200bp | 0.27% |
| –CR1: | 20 | 19115bp | 0.01% | 20 | 19115bp | 0.01% |
| –I: | 192 | 19925bp | 0.01% | 193 | 19939bp | 0.01% |
| –I-Jockey: | 9 | 4831bp | 0.00% | 9 | 4831bp | 0.00% |
| –I-Nimb: | 8 | 2342bp | 0.00% | 8 | 2342bp | 0.00% |
| –Jockey: | 12 | 1190bp | 0.00% | 12 | 1190bp | 0.00% |
| –L1-Tx1: | 4 | 629bp | 0.00% | 4 | 629bp | 0.00% |
| –L2: | 91 | 6760bp | 0.00% | 91 | 6760bp | 0.00% |
| –LOA: | 17 | 1255bp | 0.00% | 17 | 1255bp | 0.00% |
| –OTHER: | 13 | 3399bp | 0.00% | 13 | 3399bp | 0.00% |
| –Penelope: | 6790 | 664008bp | 0.22% | 6783 | 664048bp | 0.22% |
| –R1: | 1721 | 78389bp | 0.03% | 1721 | 78389bp | 0.03% |
| –R1-LOA: | 7 | 929bp | 0.00% | 7 | 929bp | 0.00% |
| –R2-NeSL: | 36 | 2863bp | 0.00% | 36 | 2863bp | 0.00% |
| –RTE-X: | 91 | 9335bp | 0.00% | 91 | 9335bp | 0.00% |
| **LTR:** | 18197 | 1058147bp | 0.36% | 18197 | 1058147bp | 0.36% |
| –Copia: | 1236 | 154160bp | 0.05% | 1236 | 154160bp | 0.05% |
| –DIRS: | 20 | 2470bp | 0.00% | 20 | 2470bp | 0.00% |
| –ERV1: | 7 | 1010bp | 0.00% | 7 | 1010bp | 0.00% |
| –Gypsy: | 10849 | 606715bp | 0.21% | 10849 | 606715bp | 0.21% |
| –Gypsy-Cigr: | 3 | 217bp | 0.00% | 3 | 217bp | 0.00% |
| –OTHER: | 26 | 3672bp | 0.00% | 26 | 3672bp | 0.00% |
| –Pao: | 6056 | 296097bp | 0.10% | 6056 | 296097bp | 0.10% |
| **RC:** | 875 | 61951bp | 0.02% | 877 | 62132bp | 0.02% |
| –Helitron: | 875 | 61951bp | 0.02% | 877 | 62132bp | 0.02% |
| **RC?:** | 34 | 2757bp | 0.00% | 34 | 2757bp | 0.00% |
| –Helitron: | 34 | 2757bp | 0.00% | 34 | 2757bp | 0.00% |
| **SINE:** | 36 | 4021bp | 0.00% | 36 | 4021bp | 0.00% |
| –5S-Deu-L2: | 16 | 1836bp | 0.00% | 16 | 1836bp | 0.00% |
| –ID: | 13 | 507bp | 0.00% | 13 | 507bp | 0.00% |
| –U: | 4 | 631bp | 0.00% | 4 | 631bp | 0.00% |
| –tRNA-RTE: | 3 | 1047bp | 0.00% | 3 | 1047bp | 0.00% |
| **Satellite:** | 8 | 510bp | 0.00% | 8 | 510bp | 0.00% |
| –OTHER: | 8 | 510bp | 0.00% | 8 | 510bp | 0.00% |
| **Simple:** | 2789 | 118800bp | 0.04% | 2789 | 118800bp | 0.04% |
| –repeat: | 2789 | 118800bp | 0.04% | 2789 | 118800bp | 0.04% |
| **Unknown:** | 90822 | 5172819bp | 1.75% | 90833 | 5174034bp | 1.75% |
| –OTHER: | 90822 | 5172819bp | 1.75% | 90833 | 5174034bp | 1.75% |
| **rRNA:** | 10 | 651bp | 0.00% | 10 | 651bp | 0.00% |
| –OTHER: | 10 | 651bp | 0.00% | 10 | 651bp | 0.00% |
| **tRNA:** | 24 | 813bp | 0.00% | 24 | 813bp | 0.00% |
| –OTHER: | 24 | 813bp | 0.00% | 24 | 813bp | 0.00% |

**Table S15. The proportion and detailed classification of detection results of SRC and RepeatClassifier covering the repetitive regions, which are generated by LongRepMarker using the reference genome of Gallus gallusbankiva as input.**

| | SparkRepeatClassifier | | | RepeatClassifier | | |
|---|---|---|---|---|---|---|
| **sequence: 464** | | | | **sequence: 464** | | |
| **total length: 1065365434bp** | | | | **total length: 1065365434bp** | | |
| **bases masked: 42433541 bp (3.98%)** | | | | **bases masked: 42433047 bp (3.98%)** | | |
| **Repeat Types** | **Num of elements** | **Length occupied** | **Percentage of sequence** | **Num of elements** | **Length occupied** | **Percentage of sequence** |
| **DNA:** | 3822 | 1511276bp | 0.14% | 3823 | 1511276bp | 0.14% |
| –Academ-1: | 23 | 2714bp | 0.00% | 23 | 2714bp | 0.00% |
| –CMC-EnSpm: | 78 | 25210bp | 0.00% | 78 | 25210bp | 0.00% |
| –Crypton-H: | 33 | 2518bp | 0.00% | 33 | 2518bp | 0.00% |
| –Crypton-V: | 8 | 6263bp | 0.00% | 8 | 6263bp | 0.00% |
| –Ginger: | 6 | 3423bp | 0.00% | 6 | 3423bp | 0.00% |
| –Kolobok: | 8 | 13471bp | 0.00% | 8 | 13471bp | 0.00% |
| –Kolobok-T2: | 4 | 10886bp | 0.00% | 4 | 10886bp | 0.00% |
| –MULE-MuDR: | 283 | 89603bp | 0.01% | 283 | 89603bp | 0.01% |
| –Maverick: | 262 | 114224bp | 0.01% | 262 | 114224bp | 0.01% |
| –OTHER: | 317 | 247749bp | 0.02% | 317 | 247749bp | 0.02% |
| –P: | 15 | 2610bp | 0.00% | 15 | 2610bp | 0.00% |
| –PIF-Harbinger: | 302 | 107185bp | 0.01% | 302 | 107185bp | 0.01% |
| –Sola-1: | 502 | 22630bp | 0.00% | 502 | 22630bp | 0.00% |
| –Sola-3: | 150 | 115060bp | 0.01% | 150 | 115060bp | 0.01% |
| –TcMar-Fot1: | 12 | 1689bp | 0.00% | 12 | 1689bp | 0.00% |
| –TcMar-ISRm11: | 2 | 1051bp | 0.00% | 2 | 1051bp | 0.00% |
| –TcMar-Mariner: | 808 | 403624bp | 0.04% | 809 | 403624bp | 0.04% |
| –TcMar-Tc1: | 38 | 69659bp | 0.01% | 38 | 69659bp | 0.01% |
| –TcMar-Tc2: | 26 | 16269bp | 0.00% | 26 | 16269bp | 0.00% |
| –Zisupton: | 4 | 316bp | 0.00% | 4 | 316bp | 0.00% |
| –hAT-Ac: | 390 | 70181bp | 0.01% | 390 | 70181bp | 0.01% |
| –hAT-Charlie: | 482 | 199204bp | 0.02% | 482 | 199204bp | 0.02% |
| –hAT-Pegasus: | 34 | 3633bp | 0.00% | 34 | 3633bp | 0.00% |
| –hAT-Tag1: | 32 | 25488bp | 0.00% | 32 | 25488bp | 0.00% |
| –hAT-Tip100: | 3 | 1764bp | 0.00% | 3 | 1764bp | 0.00% |
| **LINE:** | 56995 | 18851002bp | 1.77% | 57012 | 18854576bp | 1.77% |
| –CR1: | 55670 | 18534186bp | 1.74% | 55662 | 18533438bp | 1.74% |
| –CRE: | 14 | 5276bp | 0.00% | 14 | 5276bp | 0.00% |
| –Dualen: | 8 | 1107bp | 0.00% | 8 | 1107bp | 0.00% |
| –I: | 5 | 3620bp | 0.00% | 5 | 3620bp | 0.00% |
| –I-Jockey: | 38 | 12045bp | 0.00% | 38 | 12045bp | 0.00% |
| –Jockey: | 6 | 809bp | 0.00% | 6 | 809bp | 0.00% |
| –L1: | 149 | 72778bp | 0.01% | 149 | 72778bp | 0.01% |
| –L1-Tx1: | 3 | 1078bp | 0.00% | 3 | 1078bp | 0.00% |
| –L2: | 91 | 39329bp | 0.00% | 91 | 39329bp | 0.00% |
| –OTHER: | 36 | 63317bp | 0.01% | 36 | 63317bp | 0.01% |
| –Penelope: | 84 | 27397bp | 0.00% | 84 | 27397bp | 0.00% |
| –R1: | 18 | 5410bp | 0.00% | 18 | 5410bp | 0.00% |
| –R1-LOA: | 5 | 708bp | 0.00% | 5 | 708bp | 0.00% |
| –R2: | 108 | 44374bp | 0.00% | 133 | 49001bp | 0.00% |
| –RTE-BovB: | 621 | 190155bp | 0.02% | 621 | 190155bp | 0.02% |
| –RTE-RTE: | 4 | 3340bp | 0.00% | 4 | 3340bp | 0.00% |
| –RTE-X: | 20 | 806bp | 0.00% | 20 | 806bp | 0.00% |
| –Rex-Babar: | 21 | 7980bp | 0.00% | 21 | 7980bp | 0.00% |
| –Tad1: | 94 | 46982bp | 0.00% | 94 | 46982bp | 0.00% |
| **LTR:** | 43275 | 11135452bp | 1.05% | 43259 | 11161938bp | 1.05% |
| –Caulimovirus: | 2 | 144bp | 0.00% | 2 | 144bp | 0.00% |
| –Copia: | 327 | 175566bp | 0.02% | 331 | 175570bp | 0.02% |
| –DIRS: | 44 | 32002bp | 0.00% | 44 | 32002bp | 0.00% |
| –ERV: | 106 | 296823bp | 0.03% | 106 | 296823bp | 0.03% |
| –ERV1: | 10494 | 2734741bp | 0.26% | 10459 | 2730954bp | 0.26% |
| –ERVK: | 4556 | 1884644bp | 0.18% | 4556 | 1905031bp | 0.18% |
| –ERVL: | 26177 | 6443868bp | 0.60% | 26189 | 6486815bp | 0.61% |
| –Gypsy: | 1098 | 559901bp | 0.05% | 1098 | 559901bp | 0.05% |
| –Ngaro: | 118 | 82877bp | 0.01% | 118 | 82877bp | 0.01% |
| –OTHER: | 102 | 49259bp | 0.00% | 105 | 46763bp | 0.00% |
| –Pao: | 186 | 48475bp | 0.00% | 186 | 48475bp | 0.00% |
| –Viper: | 65 | 5407bp | 0.00% | 65 | 5407bp | 0.00% |
| **RC:** | 306 | 156071bp | 0.01% | 306 | 156071bp | 0.01% |
| –Helitron: | 306 | 156071bp | 0.01% | 306 | 156071bp | 0.01% |
| **SINE:** | 1305 | 106966bp | 0.01% | 1311 | 107869bp | 0.01% |
| –5S: | 1146 | 65107bp | 0.01% | 1146 | 65107bp | 0.01% |
| –5S-Deu-L2: | 47 | 5513bp | 0.00% | 47 | 5513bp | 0.00% |
| –5S-Sauria-RTE: | 33 | 3924bp | 0.00% | 33 | 3924bp | 0.00% |
| –Alu: | 4 | 672bp | 0.00% | 4 | 672bp | 0.00% |
| –ID: | 5 | 746bp | 0.00% | 11 | 1649bp | 0.00% |
| –MIR: | 8 | 2034bp | 0.00% | 8 | 2034bp | 0.00% |
| –U: | 42 | 2643bp | 0.00% | 42 | 2643bp | 0.00% |
| –tRNA: | 20 | 34709bp | 0.00% | 20 | 34709bp | 0.00% |
| **SINE?:** | 4 | 23697bp | 0.00% | 4 | 23697bp | 0.00% |
| –OTHER: | 4 | 23697bp | 0.00% | 4 | 23697bp | 0.00% |
| **Satellite:** | 52126 | 5494345bp | 0.52% | 52101 | 5493524bp | 0.52% |
| –OTHER: | 4012 | 954380bp | 0.09% | 3987 | 953559bp | 0.09% |
| –W-chromosome: | 39006 | 3152502bp | 0.30% | 39006 | 3152502bp | 0.30% |
| –macro: | 9035 | 1414666bp | 0.13% | 9035 | 1414666bp | 0.13% |
| –telomeric: | 73 | 3671bp | 0.00% | 73 | 3671bp | 0.00% |
| **Simple:** | 17471 | 1919486bp | 0.18% | 17471 | 1919486bp | 0.18% |
| –repeat: | 17471 | 1919486bp | 0.18% | 17471 | 1919486bp | 0.18% |
| **Unknown:** | 164468 | 14768863bp | 1.39% | 164461 | 14654956bp | 1.38% |
| –OTHER: | 164468 | 14768863bp | 1.39% | 164461 | 14654956bp | 1.38% |
| **rRNA:** | 120 | 44087bp | 0.00% | 120 | 44087bp | 0.00% |
| –OTHER: | 120 | 44087bp | 0.00% | 120 | 44087bp | 0.00% |
| **snRNA:** | 25 | 1699bp | 0.00% | 25 | 1699bp | 0.00% |
| –OTHER: | 25 | 1699bp | 0.00% | 25 | 1699bp | 0.00% |
| **tRNA:** | 245 | 32157bp | 0.00% | 245 | 32157bp | 0.00% |
| –OTHER: | 245 | 32157bp | 0.00% | 245 | 32157bp | 0.00% |

**Table S16. The proportion and detailed classification of detection results of SRC and RepeatClassifier covering the repetitive regions, which are generated by LongRepMarker using the reference genome of Glycine max as input.**

| | SparkRepeatClassifier | | | RepeatClassifier | | |
|---|---|---|---|---|---|---|
| **sequence: 1192** | | | | **sequence: 1192** | | |
| **total length: 979046046bp** | | | | **total length: 979046046bp** | | |
| **bases masked: 114369952 bp (11.68%)** | | | | **bases masked: 114368933 bp (11.68%)** | | |
| **Repeat Types** | **Num of elements** | **Length occupied** | **Percentage of sequence** | **Num of elements** | **Length occupied** | **Percentage of sequence** |
| **DNA:** | 106809 | 14542224bp | 1.49% | 106708 | 14519185bp | 1.48% |
| –Academ: | 5 | 459bp | 0.00% | 5 | 459bp | 0.00% |
| –CMC-EnSpm: | 38904 | 5942067bp | 0.61% | 38917 | 5949097bp | 0.61% |
| –Ginger: | 9 | 952bp | 0.00% | 9 | 952bp | 0.00% |
| –Kolobok-Hydra: | 4 | 9588bp | 0.00% | 4 | 9588bp | 0.00% |
| –MULE-MuDR: | 46960 | 4916688bp | 0.50% | 46860 | 4865460bp | 0.50% |
| –Maverick: | 23 | 27518bp | 0.00% | 23 | 27518bp | 0.00% |
| –MuLE-MuDR: | 7112 | 2351881bp | 0.24% | 7094 | 2365718bp | 0.24% |
| –OTHER: | 417 | 159727bp | 0.02% | 417 | 159727bp | 0.02% |
| –P: | 4 | 2481bp | 0.00% | 4 | 2481bp | 0.00% |
| –PIF-Harbinger: | 2649 | 605943bp | 0.06% | 2648 | 605943bp | 0.06% |
| –PIF-ISL2EU: | 4 | 1727bp | 0.00% | 4 | 1727bp | 0.00% |
| –PiggyBac: | 8 | 4081bp | 0.00% | 8 | 4081bp | 0.00% |
| –PiggyBac-X: | 20 | 25786bp | 0.00% | 20 | 25786bp | 0.00% |
| –Sola-2: | 4 | 777bp | 0.00% | 4 | 777bp | 0.00% |
| –TcMar-Mariner: | 4 | 4551bp | 0.00% | 4 | 4551bp | 0.00% |
| –TcMar-Pogo: | 639 | 34588bp | 0.00% | 639 | 34588bp | 0.00% |
| –TcMar-Sagan: | 4 | 615bp | 0.00% | 4 | 615bp | 0.00% |
| –TcMar-Stowaway: | 257 | 34581bp | 0.00% | 257 | 34581bp | 0.00% |
| –TcMar-Tc1: | 4 | 995bp | 0.00% | 4 | 995bp | 0.00% |
| –TcMar-Tc2: | 4 | 1260bp | 0.00% | 4 | 1260bp | 0.00% |
| –hAT-Ac: | 4312 | 567858bp | 0.06% | 4313 | 568032bp | 0.06% |
| –hAT-Charlie: | 47 | 72861bp | 0.01% | 47 | 72861bp | 0.01% |
| –hAT-Tag1: | 3393 | 609331bp | 0.06% | 3397 | 609331bp | 0.06% |
| –hAT-Tip100: | 2016 | 326734bp | 0.03% | 2016 | 326734bp | 0.03% |
| –hAT-hATm: | 6 | 922bp | 0.00% | 6 | 922bp | 0.00% |
| **LINE:** | 37045 | 6097857bp | 0.62% | 37047 | 6093704bp | 0.62% |
| –CR1: | 8 | 1506bp | 0.00% | 8 | 1506bp | 0.00% |
| –I: | 28 | 36121bp | 0.00% | 28 | 36121bp | 0.00% |
| –I-Jockey: | 32 | 14094bp | 0.00% | 32 | 14094bp | 0.00% |
| –L1: | 29043 | 4557691bp | 0.47% | 29045 | 4553547bp | 0.47% |
| –L1-DRE: | 3 | 980bp | 0.00% | 3 | 980bp | 0.00% |
| –L1-Tx1: | 8 | 13550bp | 0.00% | 8 | 13550bp | 0.00% |
| –L2: | 60 | 26784bp | 0.00% | 60 | 26784bp | 0.00% |
| –Penelope: | 17 | 1300bp | 0.00% | 17 | 1300bp | 0.00% |
| –R1: | 4 | 891bp | 0.00% | 4 | 891bp | 0.00% |
| –RTE-BovB: | 7834 | 1442598bp | 0.15% | 7834 | 1442589bp | 0.15% |
| –RTE-X: | 8 | 2343bp | 0.00% | 8 | 2343bp | 0.00% |
| **LTR:** | 295825 | 61894913bp | 6.32% | 295772 | 61898863bp | 6.32% |
| –Cassandra: | 101 | 2593bp | 0.00% | 101 | 2593bp | 0.00% |
| –Caulimovirus: | 11692 | 1038067bp | 0.11% | 11688 | 1037949bp | 0.11% |
| –Copia: | 110672 | 24312006bp | 2.48% | 110636 | 24310921bp | 2.48% |
| –ERV1: | 60 | 85955bp | 0.01% | 60 | 85955bp | 0.01% |
| –ERV4: | 4 | 5441bp | 0.00% | 4 | 5441bp | 0.00% |
| –ERVK: | 27 | 3588bp | 0.00% | 27 | 3588bp | 0.00% |
| –ERVL: | 4 | 1868bp | 0.00% | 4 | 1868bp | 0.00% |
| –Gypsy: | 172762 | 36628195bp | 3.74% | 172723 | 36606119bp | 3.74% |
| –Ngaro: | 8 | 1217bp | 0.00% | 8 | 1217bp | 0.00% |
| –OTHER: | 422 | 93749bp | 0.01% | 448 | 96647bp | 0.01% |
| –Pao: | 73 | 39864bp | 0.00% | 73 | 39864bp | 0.00% |
| **RC:** | 6109 | 1139535bp | 0.12% | 6101 | 1139331bp | 0.12% |
| –Helitron: | 6109 | 1139535bp | 0.12% | 6101 | 1139331bp | 0.12% |
| **RC?:** | 7 | 10011bp | 0.00% | 7 | 10011bp | 0.00% |
| –Helitron: | 7 | 10011bp | 0.00% | 7 | 10011bp | 0.00% |
| **Retroposon:** | 21 | 8073bp | 0.00% | 21 | 8073bp | 0.00% |
| –OTHER: | 21 | 8073bp | 0.00% | 21 | 8073bp | 0.00% |
| **SINE:** | 1330 | 172682bp | 0.02% | 1330 | 172682bp | 0.02% |
| –ID: | 29 | 1449bp | 0.00% | 29 | 1449bp | 0.00% |
| –OTHER: | 14 | 2977bp | 0.00% | 14 | 2977bp | 0.00% |
| –tRNA: | 16 | 5194bp | 0.00% | 16 | 5194bp | 0.00% |
| –tRNA-RTE: | 1271 | 167864bp | 0.02% | 1271 | 167864bp | 0.02% |
| **SINE?:** | 9 | 2128bp | 0.00% | 9 | 2128bp | 0.00% |
| –OTHER: | 9 | 2128bp | 0.00% | 9 | 2128bp | 0.00% |
| **Satellite:** | 1261 | 176627bp | 0.02% | 1261 | 176627bp | 0.02% |
| –OTHER: | 1253 | 166304bp | 0.02% | 1253 | 166304bp | 0.02% |
| –centromeric: | 8 | 10323bp | 0.00% | 8 | 10323bp | 0.00% |
| **Simple:** | 16713 | 731173bp | 0.07% | 16713 | 731173bp | 0.07% |
| –repeat: | 16713 | 731173bp | 0.07% | 16713 | 731173bp | 0.07% |
| **Unknown:** | 495250 | 35074975bp | 3.58% | 495379 | 35106006bp | 3.59% |
| –OTHER: | 495250 | 35074975bp | 3.58% | 495379 | 35106006bp | 3.59% |
| **rRNA:** | 28544 | 854797bp | 0.09% | 28544 | 854797bp | 0.09% |
| –OTHER: | 28544 | 854797bp | 0.09% | 28544 | 854797bp | 0.09% |
| **snRNA:** | 88 | 3651bp | 0.00% | 88 | 3651bp | 0.00% |
| –OTHER: | 88 | 3651bp | 0.00% | 88 | 3651bp | 0.00% |
| **tRNA:** | 644 | 33897bp | 0.00% | 644 | 33897bp | 0.00% |
| –OTHER: | 644 | 33897bp | 0.00% | 644 | 33897bp | 0.00% |

**Table S17. The proportion and detailed classification of detection results of SRC and RepeatClassifier covering the repetitive regions, which are generated by LongRepMarker using the reference genome of Mus musculus as input.**

| SparkRepeatClassifier | | | | RepeatClassifier | | |
|---|---|---|---|---|---|---|
| **sequence: 239** | | | | **sequence: 239** | | |
| **total length: 2818974548bp** | | | | **total length: 2818974548bp** | | |
| **bases masked: 260838424 bp (9.25%)** | | | | **bases masked: 260836575 bp (9.25%)** | | |
| **Repeat Types** | **Num of elements** | **Length occupied** | **Percentage of sequence** | **Num of elements** | **Length occupied** | **Percentage of sequence** |
| **DNA:** | 12442 | 2962978bp | 0.11% | 12428 | 2960246bp | 0.11% |
| –Academ: | 36 | 2694bp | 0.00% | 36 | 2694bp | 0.00% |
| –CMC-Chapaev: | 18 | 1331bp | 0.00% | 18 | 1331bp | 0.00% |
| –CMC-EnSpm: | 1327 | 324764bp | 0.01% | 1308 | 324003bp | 0.01% |
| –Crypton: | 3 | 7046bp | 0.00% | 3 | 7046bp | 0.00% |
| –Crypton-A: | 8 | 2275bp | 0.00% | 8 | 2275bp | 0.00% |
| –Crypton-F: | 4 | 556bp | 0.00% | 4 | 556bp | 0.00% |
| –Crypton-H: | 31 | 2916bp | 0.00% | 31 | 2916bp | 0.00% |
| –Crypton-S: | 24 | 1936bp | 0.00% | 24 | 1936bp | 0.00% |
| –Crypton-V: | 93 | 21394bp | 0.00% | 93 | 21394bp | 0.00% |
| –Dada: | 88 | 8493bp | 0.00% | 88 | 8493bp | 0.00% |
| –Ginger: | 187 | 32591bp | 0.00% | 197 | 32656bp | 0.00% |
| –IS3EU: | 88 | 17399bp | 0.00% | 88 | 17399bp | 0.00% |
| –Kolobok-Hydra: | 6 | 10035bp | 0.00% | 6 | 10035bp | 0.00% |
| –Kolobok-T2: | 322 | 40466bp | 0.00% | 322 | 40466bp | 0.00% |
| –MULE: | 12 | 1097bp | 0.00% | 12 | 1097bp | 0.00% |
| –MULE-MuDR: | 252 | 54222bp | 0.00% | 252 | 54222bp | 0.00% |
| –MULE-NOF: | 4 | 3892bp | 0.00% | 4 | 3892bp | 0.00% |
| –Maverick: | 92 | 14315bp | 0.00% | 92 | 14315bp | 0.00% |
| –MuLE-MuDR: | 48 | 5816bp | 0.00% | 48 | 5816bp | 0.00% |
| –MuLE-NOF: | 8 | 249bp | 0.00% | 8 | 249bp | 0.00% |
| –MuLE-NOF?: | 5 | 434bp | 0.00% | 5 | 434bp | 0.00% |
| –Novosib: | 37 | 7666bp | 0.00% | 37 | 7666bp | 0.00% |
| –OTHER: | 475 | 89064bp | 0.00% | 475 | 89064bp | 0.00% |
| –P: | 7 | 146bp | 0.00% | 7 | 146bp | 0.00% |
| –PIF-Harbinger: | 179 | 24980bp | 0.00% | 179 | 24980bp | 0.00% |
| –PIF-ISL2EU: | 61 | 5336bp | 0.00% | 61 | 5336bp | 0.00% |
| –PiggyBac: | 23 | 17800bp | 0.00% | 18 | 15764bp | 0.00% |
| –Sola-1: | 28 | 1870bp | 0.00% | 28 | 1870bp | 0.00% |
| –Sola-2: | 13 | 8115bp | 0.00% | 13 | 8115bp | 0.00% |
| –Sola-3: | 56 | 10163bp | 0.00% | 56 | 10163bp | 0.00% |
| –TcMar-IS885: | 4 | 718bp | 0.00% | 4 | 718bp | 0.00% |
| –TcMar-ISRm11: | 23 | 4096bp | 0.00% | 23 | 4096bp | 0.00% |
| –TcMar-Mariner: | 33 | 8551bp | 0.00% | 33 | 8551bp | 0.00% |
| –TcMar-Sagan: | 4 | 227bp | 0.00% | 4 | 227bp | 0.00% |
| –TcMar-Tc1: | 188 | 44269bp | 0.00% | 188 | 44269bp | 0.00% |
| –TcMar-Tc2: | 21 | 6560bp | 0.00% | 21 | 6560bp | 0.00% |
| –TcMar-Tigger: | 1762 | 293909bp | 0.01% | 1762 | 293909bp | 0.01% |
| –Zisupton: | 164 | 35683bp | 0.00% | 164 | 35683bp | 0.00% |
| –hAT: | 118 | 10626bp | 0.00% | 118 | 10626bp | 0.00% |
| –hAT-Ac: | 286 | 381717bp | 0.01% | 286 | 381717bp | 0.01% |
| –hAT-Blackjack: | 54 | 5169bp | 0.00% | 54 | 5169bp | 0.00% |
| –hAT-Charlie: | 5970 | 1415986bp | 0.05% | 5970 | 1415986bp | 0.05% |
| –hAT-Tag1: | 16 | 2181bp | 0.00% | 16 | 2181bp | 0.00% |
| –hAT-Tip100: | 195 | 67198bp | 0.00% | 195 | 67198bp | 0.00% |
| –hAT-hATw: | 69 | 6239bp | 0.00% | 69 | 6239bp | 0.00% |
| **LINE:** | 575223 | 130879619bp | 4.64% | 575238 | 130498795bp | 4.63% |
| –CR1: | 84 | 94797bp | 0.00% | 84 | 94797bp | 0.00% |
| –CRE-Ambal: | 11 | 6172bp | 0.00% | 11 | 6172bp | 0.00% |
| –I: | 39 | 66907bp | 0.00% | 39 | 66907bp | 0.00% |
| –I-Jockey: | 132 | 47906bp | 0.00% | 132 | 47906bp | 0.00% |
| –Jockey: | 17 | 2700bp | 0.00% | 17 | 2700bp | 0.00% |
| –L1: | 573389 | 130388803bp | 4.63% | 573412 | 130008161bp | 4.61% |
| –L1-Tx1: | 175 | 80576bp | 0.00% | 175 | 80576bp | 0.00% |
| –L2: | 476 | 102990bp | 0.00% | 476 | 102990bp | 0.00% |
| –LOA: | 3 | 253bp | 0.00% | 3 | 253bp | 0.00% |
| –OTHER: | 15 | 46447bp | 0.00% | 15 | 46447bp | 0.00% |
| –Penelope: | 48 | 14272bp | 0.00% | 48 | 14272bp | 0.00% |
| –R1: | 79 | 28939bp | 0.00% | 79 | 28939bp | 0.00% |
| –R2: | 62 | 48509bp | 0.00% | 54 | 47314bp | 0.00% |
| –R2-NeSL: | 8 | 2836bp | 0.00% | 8 | 2836bp | 0.00% |
| –RTE-BovB: | 607 | 155230bp | 0.01% | 607 | 155230bp | 0.01% |
| –RTE-RTE: | 8 | 2433bp | 0.00% | 8 | 2433bp | 0.00% |
| –RTE-X: | 66 | 12053bp | 0.00% | 66 | 12053bp | 0.00% |
| –Rex-Babar: | 4 | 938bp | 0.00% | 4 | 938bp | 0.00% |
| **LTR:** | 522898 | 79586508bp | 2.82% | 522837 | 79630913bp | 2.82% |
| –Caulimovirus: | 82 | 13899bp | 0.00% | 82 | 13899bp | 0.00% |
| –Copia: | 991 | 277753bp | 0.01% | 998 | 278011bp | 0.01% |
| –DIRS: | 86 | 23958bp | 0.00% | 86 | 23958bp | 0.00% |
| –ERV: | 749 | 1443253bp | 0.05% | 757 | 1446161bp | 0.05% |
| –ERV1: | 122353 | 16628052bp | 0.59% | 122350 | 16784676bp | 0.60% |
| –ERVK: | 282465 | 43357961bp | 1.54% | 282396 | 43375716bp | 1.54% |
| –ERVL: | 25836 | 8286233bp | 0.29% | 25838 | 8312011bp | 0.29% |
| –ERVL-MaLR: | 86571 | 12498839bp | 0.44% | 86561 | 12448683bp | 0.44% |
| –Gypsy: | 3149 | 591138bp | 0.02% | 3153 | 592216bp | 0.02% |
| –Lenti: | 19 | 842bp | 0.00% | 19 | 842bp | 0.00% |
| –Ngaro: | 19 | 2384bp | 0.00% | 19 | 2384bp | 0.00% |
| –OTHER: | 119 | 26437bp | 0.00% | 119 | 26437bp | 0.00% |
| –Pao: | 429 | 114264bp | 0.00% | 429 | 114264bp | 0.00% |
| –Viper: | 30 | 6470bp | 0.00% | 30 | 6470bp | 0.00% |
| **Other:** | 4696 | 698096bp | 0.02% | 4696 | 698096bp | 0.02% |
| –OTHER: | 4696 | 698096bp | 0.02% | 4696 | 698096bp | 0.02% |
| **RC:** | 385 | 46054bp | 0.00% | 385 | 46054bp | 0.00% |
| –Helitron: | 385 | 46054bp | 0.00% | 385 | 46054bp | 0.00% |
| **Retroposon:** | 10 | 1577bp | 0.00% | 10 | 1577bp | 0.00% |
| –L1: | 6 | 563bp | 0.00% | 6 | 563bp | 0.00% |
| –SVA: | 4 | 1014bp | 0.00% | 4 | 1014bp | 0.00% |
| **SINE:** | 99855 | 33312607bp | 1.18% | 99870 | 33532368bp | 1.19% |
| –5S: | 208 | 23801bp | 0.00% | 208 | 23801bp | 0.00% |
| –5S-Deu-L2: | 3 | 1522bp | 0.00% | 3 | 1522bp | 0.00% |
| –7SL: | 98 | 31742bp | 0.00% | 98 | 31742bp | 0.00% |
| –Alu: | 61154 | 26362743bp | 0.94% | 61169 | 26709402bp | 0.95% |
| –B2: | 29260 | 6335683bp | 0.22% | 29260 | 6336412bp | 0.22% |
| –B4: | 7230 | 1633364bp | 0.06% | 7230 | 1633364bp | 0.06% |
| –ID: | 1400 | 567613bp | 0.02% | 1400 | 567613bp | 0.02% |
| –MIR: | 382 | 155299bp | 0.01% | 382 | 155299bp | 0.01% |
| –U: | 80 | 7672bp | 0.00% | 80 | 7672bp | 0.00% |
| –tRNA: | 18 | 11026bp | 0.00% | 18 | 11026bp | 0.00% |
| –tRNA-7SL: | 18 | 2263bp | 0.00% | 18 | 2263bp | 0.00% |
| –tRNA-Core-L2: | 4 | 866bp | 0.00% | 4 | 866bp | 0.00% |
| **Satellite:** | 14413 | 2699863bp | 0.10% | 14417 | 2699980bp | 0.10% |
| –5S: | 4 | 1078bp | 0.00% | 0 | 0bp | 0.00% |
| –OTHER: | 14409 | 2698785bp | 0.10% | 14417 | 2699980bp | 0.10% |
| **Simple:** | 77612 | 3656460bp | 0.13% | 77612 | 3656460bp | 0.13% |
| –repeat: | 77612 | 3656460bp | 0.13% | 77612 | 3656460bp | 0.13% |
| **Unknown:** | 967058 | 60613644bp | 2.15% | 967084 | 60460592bp | 2.14% |
| –OTHER: | 964526 | 60438386bp | 2.14% | 964552 | 60285334bp | 2.14% |
| –Y-chromosome: | 2532 | 227820bp | 0.01% | 2532 | 227820bp | 0.01% |
| **rRNA:** | 204 | 11801bp | 0.00% | 204 | 11801bp | 0.00% |
| –OTHER: | 204 | 11801bp | 0.00% | 204 | 11801bp | 0.00% |
| **scRNA:** | 13 | 5519bp | 0.00% | 13 | 5519bp | 0.00% |
| –OTHER: | 13 | 5519bp | 0.00% | 13 | 5519bp | 0.00% |
| **snRNA:** | 1244 | 41044bp | 0.00% | 1244 | 41044bp | 0.00% |
| –OTHER: | 1244 | 41044bp | 0.00% | 1244 | 41044bp | 0.00% |
| **tRNA:** | 275 | 19033bp | 0.00% | 275 | 19033bp | 0.00% |
| –OTHER: | 275 | 19033bp | 0.00% | 275 | 19033bp | 0.00% |

**Table S18. The proportion and detailed classification of detection results of SRC and RepeatClassifier covering the repetitive regions, which are generated by LongRepMarker using the reference genome of Homo sapiens as input.**

| | SRC | | | RepeatClassifier | | |
|---|---|---|---|---|---|---|
| **sequence: 455** | | | | **sequence: 455** | | |
| **total length: 3209286105bp** | | | | **total length: 3209286105bp** | | |
| **bases masked: 246167245 bp (7.67%)** | | | | **bases masked: 246166512 bp (7.67%)** | | |
| **Repeat Types** | **Num of elements** | **Length occupied** | **Percentage of sequence** | **Num of elements** | **Length occupied** | **Percentage of sequence** |
| **DNA:** | 42345 | 5122539bp | 0.16% | 42348 | 5133887bp | 0.16% |
| –Academ-1: | 197 | 120305bp | 0.00% | 197 | 120305bp | 0.00% |
| –CMC-EnSpm: | 2049 | 179617bp | 0.01% | 2049 | 179617bp | 0.01% |
| –Crypton-A: | 25 | 5146bp | 0.00% | 25 | 5146bp | 0.00% |
| –Crypton-H: | 16 | 1216bp | 0.00% | 16 | 1216bp | 0.00% |
| –Crypton-S: | 5 | 2162bp | 0.00% | 5 | 2162bp | 0.00% |
| –Crypton-V: | 6 | 438bp | 0.00% | 6 | 438bp | 0.00% |
| –Dada: | 50 | 14384bp | 0.00% | 50 | 14384bp | 0.00% |
| –Ginger: | 138 | 13870bp | 0.00% | 138 | 13870bp | 0.00% |
| –IS3EU: | 18 | 4983bp | 0.00% | 18 | 4983bp | 0.00% |
| –Kolobok: | 77 | 14315bp | 0.00% | 77 | 14315bp | 0.00% |
| –Kolobok-T2: | 51 | 8646bp | 0.00% | 51 | 8646bp | 0.00% |
| –MULE-MuDR: | 334 | 45856bp | 0.00% | 334 | 45856bp | 0.00% |
| –MULE-NOF: | 10 | 2764bp | 0.00% | 10 | 2764bp | 0.00% |
| –Maverick: | 58 | 7480bp | 0.00% | 58 | 7480bp | 0.00% |
| –MuLE-MuDR: | 16 | 802bp | 0.00% | 16 | 802bp | 0.00% |
| –MuLE-NOF: | 9 | 950bp | 0.00% | 9 | 950bp | 0.00% |
| –Novosib: | 26 | 8595bp | 0.00% | 26 | 8595bp | 0.00% |
| –OTHER: | 97 | 40966bp | 0.00% | 97 | 40966bp | 0.00% |
| –P: | 10 | 841bp | 0.00% | 10 | 841bp | 0.00% |
| –PIF-Harbinger: | 34 | 10053bp | 0.00% | 34 | 10053bp | 0.00% |
| –PIF-Spy: | 4 | 226bp | 0.00% | 4 | 226bp | 0.00% |
| –PiggyBac: | 681 | 40472bp | 0.00% | 681 | 40472bp | 0.00% |
| –PiggyBac-X: | 3 | 180bp | 0.00% | 3 | 180bp | 0.00% |
| –Sola: | 3 | 143bp | 0.00% | 3 | 143bp | 0.00% |
| –Sola-1: | 12 | 394bp | 0.00% | 12 | 394bp | 0.00% |
| –Sola-2: | 18 | 978bp | 0.00% | 18 | 978bp | 0.00% |
| –Sola-3: | 29 | 10311bp | 0.00% | 29 | 10311bp | 0.00% |
| –TcMar-Fot1: | 14 | 2050bp | 0.00% | 14 | 2050bp | 0.00% |
| –TcMar-ISRm11: | 24 | 6061bp | 0.00% | 24 | 6061bp | 0.00% |
| –TcMar-Mariner: | 4232 | 243778bp | 0.01% | 4230 | 243754bp | 0.01% |
| –TcMar-Tc1: | 51 | 24889bp | 0.00% | 51 | 24889bp | 0.00% |
| –TcMar-Tc2: | 74 | 9687bp | 0.00% | 74 | 9687bp | 0.00% |
| –TcMar-Tigger: | 25728 | 1744247bp | 0.05% | 25752 | 1756174bp | 0.05% |
| –Zisupton: | 313 | 195919bp | 0.01% | 313 | 195919bp | 0.01% |
| –hAT: | 35 | 21304bp | 0.00% | 35 | 21304bp | 0.00% |
| –hAT-Ac: | 68 | 25503bp | 0.00% | 60 | 25179bp | 0.00% |
| –hAT-Blackjack: | 168 | 26762bp | 0.00% | 168 | 26762bp | 0.00% |
| –hAT-Charlie: | 6922 | 1939181bp | 0.06% | 6911 | 1938950bp | 0.06% |
| –hAT-Pegasus: | 8 | 353bp | 0.00% | 8 | 353bp | 0.00% |
| –hAT-Tag1: | 20 | 7131bp | 0.00% | 20 | 7131bp | 0.00% |
| –hAT-Tip100: | 702 | 357294bp | 0.01% | 702 | 357294bp | 0.01% |
| –hAT-Tip100?: | 4 | 1312bp | 0.00% | 4 | 1312bp | 0.00% |
| –hAT-hATm: | 6 | 2414bp | 0.00% | 6 | 2414bp | 0.00% |
| **LINE:** | 253482 | 116091245bp | 3.62% | 253551 | 116825261bp | 3.64% |
| –CR1: | 307 | 211040bp | 0.01% | 307 | 211040bp | 0.01% |
| –CRE: | 8 | 6604bp | 0.00% | 8 | 6604bp | 0.00% |
| –I: | 22 | 15578bp | 0.00% | 22 | 15578bp | 0.00% |
| –I-Jockey: | 8 | 373bp | 0.00% | 8 | 373bp | 0.00% |
| –Jockey: | 8 | 749bp | 0.00% | 8 | 749bp | 0.00% |
| –L1: | 249969 | 113923085bp | 3.55% | 250045 | 114823505bp | 3.58% |
| –L1-Tx1: | 78 | 27322bp | 0.00% | 78 | 27322bp | 0.00% |
| –L2: | 2597 | 1413646bp | 0.04% | 2597 | 1413646bp | 0.04% |
| –OTHER: | 134 | 1938577bp | 0.06% | 134 | 1938577bp | 0.06% |
| –Penelope: | 18 | 4355bp | 0.00% | 18 | 4355bp | 0.00% |
| –R1: | 75 | 49254bp | 0.00% | 75 | 49254bp | 0.00% |
| –R1-LOA: | 4 | 1556bp | 0.00% | 4 | 1556bp | 0.00% |
| –R2: | 37 | 15809bp | 0.00% | 30 | 14878bp | 0.00% |
| –RTE-BovB: | 137 | 33397bp | 0.00% | 137 | 33397bp | 0.00% |
| –RTE-X: | 68 | 25859bp | 0.00% | 68 | 25859bp | 0.00% |
| –Rex-Babar: | 12 | 2544bp | 0.00% | 12 | 2544bp | 0.00% |
| **LTR:** | 233948 | 21644832bp | 0.67% | 233960 | 21651531bp | 0.67% |
| –Caulimovirus: | 15 | 3159bp | 0.00% | 15 | 3159bp | 0.00% |
| –Copia: | 274 | 82430bp | 0.00% | 312 | 85791bp | 0.00% |
| –DIRS: | 83 | 84934bp | 0.00% | 83 | 84934bp | 0.00% |
| –ERV: | 51 | 9043bp | 0.00% | 49 | 8625bp | 0.00% |
| –ERV1: | 137547 | 9924078bp | 0.31% | 137540 | 9928300bp | 0.31% |
| –ERV4: | 12 | 13385bp | 0.00% | 12 | 13385bp | 0.00% |
| –ERVK: | 27750 | 3174763bp | 0.10% | 27757 | 3177333bp | 0.10% |
| –ERVL: | 30329 | 4440975bp | 0.14% | 30267 | 4440805bp | 0.14% |
| –ERVL-MaLR: | 36245 | 3745971bp | 0.12% | 36283 | 3742126bp | 0.12% |
| –Gypsy: | 1427 | 430273bp | 0.01% | 1427 | 430273bp | 0.01% |
| –Ngaro: | 18 | 7415bp | 0.00% | 18 | 7415bp | 0.00% |
| –OTHER: | 139 | 22011bp | 0.00% | 139 | 22011bp | 0.00% |
| –Pao: | 50 | 9434bp | 0.00% | 50 | 9434bp | 0.00% |
| –Viper: | 8 | 768bp | 0.00% | 8 | 768bp | 0.00% |
| **Other:** | 31 | 4060bp | 0.00% | 31 | 4060bp | 0.00% |
| –OTHER: | 27 | 1457bp | 0.00% | 27 | 1457bp | 0.00% |
| –subtelomeric: | 4 | 2603bp | 0.00% | 4 | 2603bp | 0.00% |
| **RC:** | 232 | 66283bp | 0.00% | 232 | 66283bp | 0.00% |
| –Helitron: | 232 | 66283bp | 0.00% | 232 | 66283bp | 0.00% |
| **RNA:** | 59 | 2808bp | 0.00% | 59 | 2808bp | 0.00% |
| –OTHER: | 59 | 2808bp | 0.00% | 59 | 2808bp | 0.00% |
| **Retroposon:** | 3377 | 1270842bp | 0.04% | 3377 | 1270884bp | 0.04% |
| –SVA: | 3377 | 1270842bp | 0.04% | 3377 | 1270884bp | 0.04% |
| **SINE:** | 129729 | 75310083bp | 2.35% | 129631 | 74868386bp | 2.33% |
| –5S: | 89 | 26210bp | 0.00% | 89 | 26210bp | 0.00% |
| –Alu: | 126088 | 74205313bp | 2.31% | 125990 | 73723264bp | 2.30% |
| –MIR: | 3456 | 1439502bp | 0.04% | 3456 | 1439502bp | 0.04% |
| –U: | 62 | 14456bp | 0.00% | 62 | 14456bp | 0.00% |
| –tRNA-7SL: | 16 | 1411bp | 0.00% | 16 | 1411bp | 0.00% |
| –tRNA-Core-L2: | 4 | 184bp | 0.00% | 4 | 184bp | 0.00% |
| –tRNA-Meta: | 10 | 1898bp | 0.00% | 10 | 1898bp | 0.00% |
| –tRNA-RTE: | 4 | 179bp | 0.00% | 4 | 179bp | 0.00% |
| **SINE?:** | 16 | 21177bp | 0.00% | 16 | 21177bp | 0.00% |
| –OTHER: | 16 | 21177bp | 0.00% | 16 | 21177bp | 0.00% |
| **Satellite:** | 1339490 | 51375051bp | 1.60% | 1339459 | 51375863bp | 1.60% |
| –OTHER: | 8157 | 1392631bp | 0.04% | 8164 | 1393562bp | 0.04% |
| –Y-chromosome: | 321517 | 24008707bp | 0.75% | 321484 | 24005856bp | 0.75% |
| –acromeric: | 203 | 91137bp | 0.00% | 203 | 91137bp | 0.00% |
| –centromeric: | 1008827 | 43052149bp | 1.34% | 1008822 | 43052590bp | 1.34% |
| –telomeric: | 786 | 192714bp | 0.01% | 786 | 192714bp | 0.01% |
| **Simple:** | 19483 | 1534323bp | 0.05% | 19483 | 1534323bp | 0.05% |
| –repeat: | 19483 | 1534323bp | 0.05% | 19483 | 1534323bp | 0.05% |
| **Unknown:** | 158348 | 38949784bp | 1.21% | 158386 | 39286803bp | 1.22% |
| –OTHER: | 158348 | 38949784bp | 1.21% | 158386 | 39286803bp | 1.22% |
| **rRNA:** | 256 | 98971bp | 0.00% | 256 | 98971bp | 0.00% |
| –OTHER: | 256 | 98971bp | 0.00% | 256 | 98971bp | 0.00% |
| **scRNA:** | 35 | 3213bp | 0.00% | 35 | 3213bp | 0.00% |
| –OTHER: | 35 | 3213bp | 0.00% | 35 | 3213bp | 0.00% |
| **snRNA:** | 140 | 46208bp | 0.00% | 140 | 46208bp | 0.00% |
| –OTHER: | 140 | 46208bp | 0.00% | 140 | 46208bp | 0.00% |
| **tRNA:** | 235 | 19643bp | 0.00% | 235 | 19643bp | 0.00% |
| –OTHER: | 235 | 19643bp | 0.00% | 235 | 19643bp | 0.00% |

**Table S19. The proportion and detailed classification of detection results of SRC and RepeatClassifier covering the repetitive regions, which are generated by RepeatScout using the reference genome of Picea abies as input.**

| | SparkRepeatClassifier | | | RepeatClassifier | | |
|---|---|---|---|---|---|---|
| **sequence: 11340369** | | | | **sequence: 11340369** | | |
| **total length: 11961396284bp** | | | | **total length: 11961396284bp** | | |
| **bases masked: 1610279537 bp (13.46%)** | | | | **bases masked: 1610114420 bp (13.46%)** | | |
| **Repeat Types** | **Num of elements** | **Length occupied** | **Percentage of sequence** | **Num of elements** | **Length occupied** | **Percentage of sequence** |
| **ARTEFACT:** | 28 | 8623bp | 0.00% | 28 | 8623bp | 0.00% |
| –OTHER: | 28 | 8623bp | 0.00% | 28 | 8623bp | 0.00% |
| **DNA:** | 201492 | 23358088bp | 0.20% | 201627 | 23360691bp | 0.20% |
| –Academ-1: | 353 | 73666bp | 0.00% | 353 | 73517bp | 0.00% |
| –Academ-2: | 44 | 6508bp | 0.00% | 44 | 6608bp | 0.00% |
| –CMC-Chapaev: | 40 | 13018bp | 0.00% | 40 | 13018bp | 0.00% |
| –CMC-Chapaev-3: | 145 | 26281bp | 0.00% | 145 | 26353bp | 0.00% |
| –CMC-EnSpm: | 76500 | 10030634bp | 0.08% | 76711 | 10036283bp | 0.08% |
| –CMC-Mirage: | 17 | 6581bp | 0.00% | 17 | 6581bp | 0.00% |
| –CMC-Transib: | 67 | 10436bp | 0.00% | 67 | 10440bp | 0.00% |
| –Crypton-H: | 545 | 97902bp | 0.00% | 545 | 98719bp | 0.00% |
| –Crypton-R: | 83 | 10997bp | 0.00% | 83 | 10997bp | 0.00% |
| –Crypton-V: | 124 | 7168bp | 0.00% | 124 | 7146bp | 0.00% |
| –Dada: | 2238 | 398411bp | 0.00% | 2239 | 399235bp | 0.00% |
| –Ginger: | 268 | 39807bp | 0.00% | 356 | 56005bp | 0.00% |
| –IS3EU: | 1479 | 123389bp | 0.00% | 1478 | 123594bp | 0.00% |
| –Kolobok-E: | 48 | 5236bp | 0.00% | 48 | 5234bp | 0.00% |
| –Kolobok-Hydra: | 79 | 13473bp | 0.00% | 79 | 13229bp | 0.00% |
| –Kolobok-T2: | 256 | 53630bp | 0.00% | 256 | 53627bp | 0.00% |
| –MULE-MuDR: | 3476 | 578578bp | 0.00% | 3493 | 580079bp | 0.00% |
| –MULE-NOF: | 101 | 12617bp | 0.00% | 101 | 12586bp | 0.00% |
| –Maverick: | 993 | 215166bp | 0.00% | 1028 | 180913bp | 0.00% |
| –Merlin: | 103 | 16443bp | 0.00% | 103 | 16444bp | 0.00% |
| –MuLE-MuDR: | 3252 | 972754bp | 0.01% | 3284 | 1083180bp | 0.01% |
| –OTHER: | 4901 | 656996bp | 0.01% | 4785 | 626986bp | 0.01% |
| –P: | 2081 | 302549bp | 0.00% | 2081 | 302998bp | 0.00% |
| –PIF-HarbS: | 39 | 13837bp | 0.00% | 39 | 13845bp | 0.00% |
| –PIF-Harbinger: | 1499 | 312705bp | 0.00% | 1434 | 270833bp | 0.00% |
| –PIF-ISL2EU: | 59 | 18174bp | 0.00% | 59 | 18174bp | 0.00% |
| –PIF-Spy: | 111 | 24179bp | 0.00% | 111 | 24179bp | 0.00% |
| –PiggyBac: | 621 | 179706bp | 0.00% | 616 | 176408bp | 0.00% |
| –PiggyBac-X: | 48 | 17338bp | 0.00% | 48 | 17338bp | 0.00% |
| –Sola: | 9060 | 1832226bp | 0.02% | 9680 | 1906731bp | 0.02% |
| –Sola-1: | 1740 | 508354bp | 0.00% | 1308 | 463416bp | 0.00% |
| –Sola-2: | 1851 | 665131bp | 0.01% | 1693 | 541262bp | 0.00% |
| –TcMar: | 100 | 31857bp | 0.00% | 100 | 31857bp | 0.00% |
| –TcMar-Ant1: | 1176 | 182464bp | 0.00% | 1176 | 182301bp | 0.00% |
| –TcMar-Fot1: | 66 | 20006bp | 0.00% | 20 | 15008bp | 0.00% |
| –TcMar-ISRm11: | 72 | 16088bp | 0.00% | 72 | 16088bp | 0.00% |
| –TcMar-Mariner: | 87 | 29737bp | 0.00% | 87 | 29784bp | 0.00% |
| –TcMar-Pogo: | 107 | 42707bp | 0.00% | 107 | 42707bp | 0.00% |
| –TcMar-Sagan: | 100 | 15069bp | 0.00% | 100 | 15042bp | 0.00% |
| –TcMar-Tc1: | 337 | 109568bp | 0.00% | 337 | 109556bp | 0.00% |
| –TcMar-Tc2: | 33 | 6157bp | 0.00% | 33 | 6157bp | 0.00% |
| –TcMar-Tc4: | 132 | 18398bp | 0.00% | 132 | 18398bp | 0.00% |
| –TcMar-Tigger: | 30 | 4665bp | 0.00% | 30 | 4665bp | 0.00% |
| –Zisupton: | 39 | 19629bp | 0.00% | 39 | 19629bp | 0.00% |
| –hAT: | 42 | 3154bp | 0.00% | 42 | 3163bp | 0.00% |
| –hAT-Ac: | 2597 | 346683bp | 0.00% | 2522 | 289789bp | 0.00% |
| –hAT-Blackjack: | 51 | 51420bp | 0.00% | 51 | 51420bp | 0.00% |
| –hAT-Charlie: | 8883 | 784238bp | 0.01% | 8875 | 781681bp | 0.01% |
| –hAT-Tag1: | 71588 | 4688831bp | 0.04% | 71596 | 4693409bp | 0.04% |
| –hAT-Tip100: | 2898 | 481502bp | 0.00% | 2969 | 497948bp | 0.00% |
| –hAT-hAT1: | 42 | 18619bp | 0.00% | 0 | 0bp | 0.00% |
| –hAT-hAT5: | 3 | 117bp | 0.00% | 3 | 117bp | 0.00% |
| –hAT-hATm: | 276 | 28770bp | 0.00% | 276 | 28582bp | 0.00% |
| –hAT-hATw: | 135 | 62983bp | 0.00% | 135 | 62983bp | 0.00% |
| –hAT-hobo: | 477 | 338439bp | 0.00% | 477 | 336812bp | 0.00% |
| **LINE:** | 271849 | 32920281bp | 0.28% | 272803 | 33004764bp | 0.28% |
| –CR1: | 2555 | 700766bp | 0.01% | 2586 | 739254bp | 0.01% |
| –CR1-Zenon: | 76 | 9487bp | 0.00% | 0 | 0bp | 0.00% |
| –Dong-R4: | 190 | 17512bp | 0.00% | 190 | 17555bp | 0.00% |
| –I: | 4213 | 908450bp | 0.01% | 4257 | 903653bp | 0.01% |
| –I-Jockey: | 944 | 114465bp | 0.00% | 916 | 113242bp | 0.00% |
| –I-Nimb: | 227 | 43437bp | 0.00% | 247 | 49772bp | 0.00% |
| –Jockey: | 674 | 150317bp | 0.00% | 656 | 132813bp | 0.00% |
| –L1: | 196538 | 22863853bp | 0.19% | 196592 | 22858879bp | 0.19% |
| –L1-DRE: | 29 | 2968bp | 0.00% | 29 | 2968bp | 0.00% |
| –L1-Tx1: | 4303 | 827635bp | 0.01% | 4607 | 877026bp | 0.01% |
| –L2: | 4661 | 956976bp | 0.01% | 4298 | 850148bp | 0.01% |
| –OTHER: | 3180 | 916033bp | 0.01% | 3441 | 958186bp | 0.01% |
| –Penelope: | 13890 | 1410346bp | 0.01% | 13888 | 1414807bp | 0.01% |
| –Proto2: | 783 | 156328bp | 0.00% | 783 | 155469bp | 0.00% |
| –R1: | 657 | 110283bp | 0.00% | 1080 | 221485bp | 0.00% |
| –R1-LOA: | 139 | 17429bp | 0.00% | 142 | 17553bp | 0.00% |
| –R2: | 228 | 43365bp | 0.00% | 251 | 48308bp | 0.00% |
| –R2-Hero: | 56 | 15252bp | 0.00% | 56 | 15252bp | 0.00% |
| –R2-NeSL: | 2334 | 456849bp | 0.00% | 1897 | 356797bp | 0.00% |
| –RTE-BovB: | 3653 | 577102bp | 0.00% | 3473 | 601819bp | 0.01% |
| –RTE-RTE: | 154 | 15554bp | 0.00% | 102 | 14189bp | 0.00% |
| –RTE-X: | 30326 | 3987762bp | 0.03% | 31246 | 4007848bp | 0.03% |
| –Rex-Babar: | 44 | 2234bp | 0.00% | 59 | 8211bp | 0.00% |
| –Tad1: | 1289 | 304549bp | 0.00% | 1228 | 283020bp | 0.00% |
| –Tad1?: | 706 | 110817bp | 0.00% | 779 | 115704bp | 0.00% |
| **LTR:** | 11039203 | 1042914597bp | 8.72% | 11036954 | 1042517160bp | 8.72% |
| –Bhikhari: | 49 | 7637bp | 0.00% | 49 | 7637bp | 0.00% |
| –Cassandra: | 8370 | 612272bp | 0.01% | 8370 | 611856bp | 0.01% |
| –Caulimovirus: | 6443 | 1177040bp | 0.01% | 6678 | 1206186bp | 0.01% |
| –Copia: | 2439969 | 226243200bp | 1.89% | 2438400 | 226112241bp | 1.89% |
| –DIRS: | 1129 | 271120bp | 0.00% | 1151 | 270191bp | 0.00% |
| –ERV: | 85 | 13107bp | 0.00% | 37 | 4383bp | 0.00% |
| –ERV-Lenti: | 0 | 0bp | 0.00% | 28 | 11985bp | 0.00% |
| –ERV1: | 3391 | 616602bp | 0.01% | 3149 | 603574bp | 0.01% |
| –ERV4: | 79 | 7820bp | 0.00% | 79 | 7820bp | 0.00% |
| –ERVK: | 4460 | 807661bp | 0.01% | 4017 | 795067bp | 0.01% |
| –ERVL: | 343 | 93808bp | 0.00% | 343 | 93808bp | 0.00% |
| –ERVL-MaLR: | 17 | 4716bp | 0.00% | 17 | 4716bp | 0.00% |
| –Foamy: | 56 | 2920bp | 0.00% | 129 | 4688bp | 0.00% |
| –Gypsy: | 8552729 | 820141182bp | 6.86% | 8552965 | 820256548bp | 6.86% |
| –Gypsy-Cigr: | 174 | 20746bp | 0.00% | 321 | 36413bp | 0.00% |
| –Lenti: | 423 | 76296bp | 0.00% | 301 | 39062bp | 0.00% |
| –Ngaro: | 193 | 18212bp | 0.00% | 193 | 18214bp | 0.00% |
| –OTHER: | 16381 | 4088896bp | 0.03% | 15641 | 3521834bp | 0.03% |
| –Pao: | 4912 | 1113609bp | 0.01% | 5086 | 1067664bp | 0.01% |
| **RC:** | 35678 | 4113280bp | 0.03% | 36143 | 4205863bp | 0.04% |
| –Helitron: | 35678 | 4113280bp | 0.03% | 36143 | 4205863bp | 0.04% |
| **Retroposon:** | 149 | 24772bp | 0.00% | 149 | 24683bp | 0.00% |
| –OTHER: | 149 | 24772bp | 0.00% | 149 | 24683bp | 0.00% |
| **SINE:** | 503 | 94572bp | 0.00% | 503 | 94623bp | 0.00% |
| –7SL: | 36 | 2740bp | 0.00% | 36 | 2740bp | 0.00% |
| –ID: | 154 | 2697bp | 0.00% | 154 | 2748bp | 0.00% |
| –U: | 113 | 17067bp | 0.00% | 113 | 17067bp | 0.00% |
| –tRNA: | 66 | 4537bp | 0.00% | 66 | 4537bp | 0.00% |
| –tRNA-V: | 134 | 67531bp | 0.00% | 134 | 67531bp | 0.00% |
| **Satellite:** | 31225 | 915070bp | 0.01% | 31231 | 915797bp | 0.01% |
| –5S: | 544 | 47091bp | 0.00% | 544 | 46788bp | 0.00% |
| –OTHER: | 30642 | 844436bp | 0.01% | 30648 | 845466bp | 0.01% |
| –centromeric: | 39 | 23543bp | 0.00% | 39 | 23543bp | 0.00% |
| **Simple:** | 221078 | 7556589bp | 0.06% | 221052 | 7538899bp | 0.06% |
| –repeat: | 221078 | 7556589bp | 0.06% | 221052 | 7538899bp | 0.06% |
| **Unknown:** | 10082618 | 616599349bp | 5.15% | 10081629 | 616261277bp | 5.15% |
| –OTHER: | 10082618 | 616599349bp | 5.15% | 10081629 | 616261277bp | 5.15% |
| **rRNA:** | 7226 | 302432bp | 0.00% | 7226 | 302516bp | 0.00% |
| –OTHER: | 7226 | 302432bp | 0.00% | 7226 | 302516bp | 0.00% |
| **snRNA:** | 116 | 5405bp | 0.00% | 116 | 5405bp | 0.00% |
| –OTHER: | 116 | 5405bp | 0.00% | 116 | 5405bp | 0.00% |
| **tRNA:** | 2102 | 90891bp | 0.00% | 2102 | 90755bp | 0.00% |
| –OTHER: | 2102 | 90891bp | 0.00% | 2102 | 90755bp | 0.00% |

**Table S20. The proportion and detailed classification of detection results of SRC and RepeatClassifier covering the repetitive regions, which are generated by RepeatScout using the reference genome of Mus musculus as input.**

| | SparkRepeatClassifier | | | RepeatClassifier | | |
|---|---|---|---|---|---|---|
| | **sequence: 239** | | | **sequence: 239** | | |
| | **total length: 2818974548bp** | | | **total length: 2818974548bp** | | |
| | **bases masked: 133720744 bp (4.74%)** | | | **bases masked: 133723208 bp (4.74%)** | | |
| **Repeat Types** | **Num of elements** | **Length occupied** | **Percentage of sequence** | **Num of elements** | **Length occupied** | **Percentage of sequence** |
| **DNA:** | 10145 | 1668081bp | 0.06% | 10093 | 1698500bp | 0.06% |
| –Academ: | 91 | 8053bp | 0.00% | 39 | 4198bp | 0.00% |
| –CMC-Chapaev: | 8 | 5216bp | 0.00% | 0 | 0bp | 0.00% |
| –CMC-Chapaev-3: | 16 | 2900bp | 0.00% | 16 | 2900bp | 0.00% |
| –CMC-EnSpm: | 280 | 106615bp | 0.00% | 280 | 106607bp | 0.00% |
| –Crypton: | 17 | 10982bp | 0.00% | 33 | 12389bp | 0.00% |
| –Dada: | 38 | 6891bp | 0.00% | 38 | 6891bp | 0.00% |
| –Ginger: | 56 | 15635bp | 0.00% | 56 | 15635bp | 0.00% |
| –IS3EU: | 67 | 18943bp | 0.00% | 67 | 18943bp | 0.00% |
| –Kolobok-Hydra: | 29 | 4557bp | 0.00% | 29 | 4557bp | 0.00% |
| –Kolobok-T2: | 12 | 3750bp | 0.00% | 12 | 3750bp | 0.00% |
| –MULE-MuDR: | 286 | 46274bp | 0.00% | 286 | 46274bp | 0.00% |
| –Maverick: | 93 | 24063bp | 0.00% | 93 | 24063bp | 0.00% |
| –MuLE-MuDR: | 35 | 4343bp | 0.00% | 35 | 4343bp | 0.00% |
| –Novosib: | 58 | 5806bp | 0.00% | 58 | 5806bp | 0.00% |
| –OTHER: | 179 | 115765bp | 0.00% | 168 | 121079bp | 0.00% |
| –PIF-Harbinger: | 253 | 39238bp | 0.00% | 253 | 39238bp | 0.00% |
| –PiggyBac: | 64 | 24028bp | 0.00% | 54 | 23944bp | 0.00% |
| –PiggyBac-X: | 11 | 1482bp | 0.00% | 11 | 1482bp | 0.00% |
| –Sola-2: | 14 | 3446bp | 0.00% | 14 | 3446bp | 0.00% |
| –Sola-3: | 28 | 2876bp | 0.00% | 28 | 2876bp | 0.00% |
| –TcMar: | 16 | 969bp | 0.00% | 16 | 969bp | 0.00% |
| –TcMar-Ant1: | 8 | 1080bp | 0.00% | 8 | 1080bp | 0.00% |
| –TcMar-ISRm11: | 12 | 4352bp | 0.00% | 12 | 4352bp | 0.00% |
| –TcMar-Tc2: | 24 | 4696bp | 0.00% | 24 | 4696bp | 0.00% |
| –TcMar-Tigger: | 265 | 52357bp | 0.00% | 265 | 52357bp | 0.00% |
| –Zisupton: | 42 | 9898bp | 0.00% | 42 | 9898bp | 0.00% |
| –hAT-Ac: | 176 | 64988bp | 0.00% | 194 | 94134bp | 0.00% |
| –hAT-Charlie: | 7768 | 1034079bp | 0.04% | 7763 | 1034125bp | 0.04% |
| –hAT-Tag1: | 72 | 18044bp | 0.00% | 72 | 18044bp | 0.00% |
| –hAT-Tip100: | 127 | 51289bp | 0.00% | 127 | 51289bp | 0.00% |
| **LINE:** | 78485 | 61591626bp | 2.18% | 78853 | 61848695bp | 2.19% |
| –CR1: | 17 | 61029bp | 0.00% | 17 | 61029bp | 0.00% |
| –CRE-Ambal: | 28 | 5980bp | 0.00% | 28 | 5980bp | 0.00% |
| –I: | 92 | 59137bp | 0.00% | 212 | 168490bp | 0.01% |
| –I-Jockey: | 92 | 16672bp | 0.00% | 92 | 16672bp | 0.00% |
| –L1: | 76927 | 61152562bp | 2.17% | 76953 | 61290687bp | 2.17% |
| –L1-Tx1: | 67 | 114319bp | 0.00% | 67 | 114319bp | 0.00% |
| –L2: | 377 | 72643bp | 0.00% | 377 | 72643bp | 0.00% |
| –R1: | 31 | 8611bp | 0.00% | 31 | 8611bp | 0.00% |
| –R2: | 21 | 10650bp | 0.00% | 243 | 31733bp | 0.00% |
| –RTE-BovB: | 829 | 248992bp | 0.01% | 829 | 249314bp | 0.01% |
| –RTE-X: | 4 | 350bp | 0.00% | 4 | 350bp | 0.00% |
| **LTR:** | 136427 | 41579312bp | 1.47% | 136551 | 41061967bp | 1.46% |
| –Copia: | 363 | 139928bp | 0.00% | 363 | 139928bp | 0.00% |
| –DIRS: | 82 | 25787bp | 0.00% | 82 | 25787bp | 0.00% |
| –ERV: | 189 | 2263584bp | 0.08% | 204 | 2275532bp | 0.08% |
| –ERV1: | 18716 | 7638964bp | 0.27% | 18593 | 7564513bp | 0.27% |
| –ERV4: | 45 | 6877bp | 0.00% | 45 | 6877bp | 0.00% |
| –ERVK: | 82238 | 21833803bp | 0.77% | 82443 | 21879723bp | 0.78% |
| –ERVL: | 7727 | 3337160bp | 0.12% | 7643 | 3308644bp | 0.12% |
| –ERVL-MaLR: | 24189 | 7298575bp | 0.26% | 24325 | 6800066bp | 0.24% |
| –Gypsy: | 2277 | 433422bp | 0.02% | 2260 | 433422bp | 0.02% |
| –OTHER: | 22 | 5674bp | 0.00% | 22 | 5674bp | 0.00% |
| –Pao: | 579 | 61107bp | 0.00% | 571 | 60107bp | 0.00% |
| **Other:** | 2418 | 166407bp | 0.01% | 2418 | 166258bp | 0.01% |
| –OTHER: | 2418 | 166407bp | 0.01% | 2418 | 166258bp | 0.01% |
| **RC:** | 384 | 54537bp | 0.00% | 390 | 54537bp | 0.00% |
| –Helitron: | 384 | 54537bp | 0.00% | 390 | 54537bp | 0.00% |
| **RNA:** | 25 | 881bp | 0.00% | 25 | 881bp | 0.00% |
| –OTHER: | 25 | 881bp | 0.00% | 25 | 881bp | 0.00% |
| **SINE:** | 63375 | 20335828bp | 0.72% | 63450 | 20385534bp | 0.72% |
| –5S: | 60 | 3916bp | 0.00% | 60 | 3994bp | 0.00% |
| –5S-Deu-L2: | 9 | 3436bp | 0.00% | 9 | 3436bp | 0.00% |
| –7SL: | 27 | 1464bp | 0.00% | 27 | 1464bp | 0.00% |
| –Alu: | 31688 | 14416083bp | 0.51% | 31704 | 14464370bp | 0.51% |
| –B2: | 24911 | 5399987bp | 0.19% | 24954 | 5399751bp | 0.19% |
| –B4: | 4719 | 1068522bp | 0.04% | 4735 | 1069484bp | 0.04% |
| –ID: | 1270 | 298677bp | 0.01% | 1270 | 298677bp | 0.01% |
| –MIR: | 452 | 150017bp | 0.01% | 452 | 150017bp | 0.01% |
| –U: | 203 | 6912bp | 0.00% | 203 | 6863bp | 0.00% |
| –tRNA: | 36 | 1620bp | 0.00% | 36 | 1620bp | 0.00% |
| **Satellite:** | 3512 | 1409616bp | 0.05% | 3153 | 1353960bp | 0.05% |
| –OTHER: | 3512 | 1409616bp | 0.05% | 3153 | 1353960bp | 0.05% |
| **Simple:** | 118608 | 4221854bp | 0.15% | 118598 | 4221405bp | 0.15% |
| –repeat: | 118608 | 4221854bp | 0.15% | 118598 | 4221405bp | 0.15% |
| **Unknown:** | 113653 | 14506163bp | 0.51% | 113522 | 14432946bp | 0.51% |
| –OTHER: | 113653 | 14506163bp | 0.51% | 113522 | 14432946bp | 0.51% |
| **rRNA:** | 157 | 9129bp | 0.00% | 157 | 9129bp | 0.00% |
| –OTHER: | 157 | 9129bp | 0.00% | 157 | 9129bp | 0.00% |
| **snRNA:** | 593 | 41432bp | 0.00% | 593 | 41433bp | 0.00% |
| –OTHER: | 593 | 41432bp | 0.00% | 593 | 41433bp | 0.00% |
| **tRNA:** | 391 | 34168bp | 0.00% | 391 | 33998bp | 0.00% |
| –OTHER: | 391 | 34168bp | 0.00% | 391 | 33998bp | 0.00% |

**Table S21. The proportion and detailed classification of detection results of SRC and RepeatClassifier covering the repetitive regions, which are generated by RepeatScout using the reference genome of Homo sapiens as input.**

| | SparkRepeatClassifier | | | RepeatClassifier | | |
|---|---|---|---|---|---|---|
| **sequence: 455** | | | | **sequence: 455** | | |
| **total length: 3209286105bp** | | | | **total length: 3209286105bp** | | |
| **bases masked: 133290370 bp (4.15%)** | | | | **bases masked: 133352006 bp (4.16%)** | | |
| **Repeat Types** | **Num of elements** | **Length occupied** | **Percentage of sequence** | **Num of elements** | **Length occupied** | **Percentage of sequence** |
| **DNA:** | 32016 | 4783695bp | 0.15% | 32095 | 4756426bp | 0.15% |
| –Academ-1: | 133 | 178620bp | 0.01% | 133 | 178620bp | 0.01% |
| –CMC-EnSpm: | 142 | 30651bp | 0.00% | 142 | 30651bp | 0.00% |
| –Crypton-V: | 35 | 14606bp | 0.00% | 35 | 14606bp | 0.00% |
| –Dada: | 6 | 615bp | 0.00% | 6 | 615bp | 0.00% |
| –IS3EU: | 31 | 5558bp | 0.00% | 31 | 5558bp | 0.00% |
| –Kolobok-T2: | 1183 | 93490bp | 0.00% | 1184 | 93490bp | 0.00% |
| –MULE-MuDR: | 428 | 76737bp | 0.00% | 434 | 77130bp | 0.00% |
| –Maverick: | 14 | 1732bp | 0.00% | 14 | 1732bp | 0.00% |
| –Merlin: | 1 | 41bp | 0.00% | 1 | 41bp | 0.00% |
| –MuLE-MuDR: | 51 | 6620bp | 0.00% | 64 | 6936bp | 0.00% |
| –OTHER: | 74 | 21960bp | 0.00% | 74 | 21960bp | 0.00% |
| –P: | 32 | 6957bp | 0.00% | 32 | 6957bp | 0.00% |
| –PIF-Harbinger: | 46 | 4239bp | 0.00% | 46 | 4239bp | 0.00% |
| –PiggyBac: | 517 | 41673bp | 0.00% | 518 | 40931bp | 0.00% |
| –PiggyBac-X: | 16 | 2088bp | 0.00% | 16 | 2088bp | 0.00% |
| –Sola-1: | 52 | 15003bp | 0.00% | 52 | 15003bp | 0.00% |
| –Sola-3: | 48 | 4239bp | 0.00% | 48 | 4239bp | 0.00% |
| –TcMar-Fot1: | 9 | 948bp | 0.00% | 9 | 948bp | 0.00% |
| –TcMar-Mariner: | 744 | 189296bp | 0.01% | 744 | 189296bp | 0.01% |
| –TcMar-Tc1: | 36 | 38040bp | 0.00% | 36 | 38040bp | 0.00% |
| –TcMar-Tc2: | 195 | 47424bp | 0.00% | 195 | 47424bp | 0.00% |
| –TcMar-Tigger: | 14481 | 1820562bp | 0.06% | 14489 | 1847063bp | 0.06% |
| –Zisupton: | 614 | 75048bp | 0.00% | 614 | 75048bp | 0.00% |
| –hAT: | 37 | 1964bp | 0.00% | 37 | 1964bp | 0.00% |
| –hAT-Ac: | 60 | 106968bp | 0.00% | 50 | 128670bp | 0.00% |
| –hAT-Blackjack: | 493 | 40597bp | 0.00% | 493 | 40597bp | 0.00% |
| –hAT-Charlie: | 11251 | 1468013bp | 0.05% | 11311 | 1392864bp | 0.04% |
| –hAT-Tag1: | 14 | 14793bp | 0.00% | 14 | 14793bp | 0.00% |
| –hAT-Tip100: | 1273 | 487371bp | 0.02% | 1273 | 487371bp | 0.02% |
| **DNA?:** | 15 | 2951bp | 0.00% | 15 | 2951bp | 0.00% |
| –hAT-Tip100: | 15 | 2951bp | 0.00% | 15 | 2951bp | 0.00% |
| **LINE:** | 96837 | 41584846bp | 1.30% | 96753 | 37439149bp | 1.17% |
| –CR1: | 303 | 229534bp | 0.01% | 303 | 229534bp | 0.01% |
| –I: | 234 | 139199bp | 0.00% | 215 | 136569bp | 0.00% |
| –I-Jockey: | 33 | 9887bp | 0.00% | 33 | 9887bp | 0.00% |
| –I-Nimb: | 6 | 870bp | 0.00% | 6 | 870bp | 0.00% |
| –L1: | 92085 | 39724226bp | 1.24% | 92148 | 35612579bp | 1.11% |
| –L1-Tx1: | 314 | 57825bp | 0.00% | 116 | 45221bp | 0.00% |
| –L2: | 3264 | 1355378bp | 0.04% | 3306 | 1278158bp | 0.04% |
| –OTHER: | 67 | 174921bp | 0.01% | 95 | 175321bp | 0.01% |
| –Penelope: | 6 | 10617bp | 0.00% | 6 | 10617bp | 0.00% |
| –R1: | 203 | 22192bp | 0.00% | 203 | 22192bp | 0.00% |
| –R2: | 22 | 1619bp | 0.00% | 22 | 1619bp | 0.00% |
| –R2-Hero: | 8 | 1394bp | 0.00% | 8 | 1394bp | 0.00% |
| –RTE-BovB: | 204 | 81103bp | 0.00% | 204 | 81103bp | 0.00% |
| –RTE-X: | 88 | 30551bp | 0.00% | 88 | 30551bp | 0.00% |
| **LTR:** | 147815 | 22075501bp | 0.69% | 147377 | 21992807bp | 0.69% |
| –Copia: | 500 | 91543bp | 0.00% | 314 | 77033bp | 0.00% |
| –DIRS: | 50 | 58539bp | 0.00% | 54 | 58601bp | 0.00% |
| –ERV: | 103 | 92592bp | 0.00% | 166 | 103236bp | 0.00% |
| –ERV1: | 68850 | 10660563bp | 0.33% | 68726 | 10635143bp | 0.33% |
| –ERVK: | 11074 | 3096862bp | 0.10% | 11052 | 3107476bp | 0.10% |
| –ERVL: | 19691 | 3920946bp | 0.12% | 19444 | 3898745bp | 0.12% |
| –ERVL-MaLR: | 45790 | 4253527bp | 0.13% | 45871 | 4160292bp | 0.13% |
| –Gypsy: | 1220 | 391359bp | 0.01% | 1213 | 391033bp | 0.01% |
| –Ngaro: | 314 | 36778bp | 0.00% | 314 | 36763bp | 0.00% |
| –OTHER: | 125 | 37855bp | 0.00% | 125 | 37855bp | 0.00% |
| –Pao: | 86 | 12348bp | 0.00% | 86 | 12348bp | 0.00% |
| –Viper: | 12 | 1624bp | 0.00% | 12 | 1624bp | 0.00% |
| **RC:** | 138 | 35797bp | 0.00% | 138 | 35797bp | 0.00% |
| –Helitron: | 138 | 35797bp | 0.00% | 138 | 35797bp | 0.00% |
| **RC?:** | 29 | 2460bp | 0.00% | 29 | 2460bp | 0.00% |
| –Helitron: | 29 | 2460bp | 0.00% | 29 | 2460bp | 0.00% |
| **RNA:** | 77 | 4050bp | 0.00% | 77 | 4050bp | 0.00% |
| –OTHER: | 77 | 4050bp | 0.00% | 77 | 4050bp | 0.00% |
| **Retroposon:** | 1955 | 540441bp | 0.02% | 1955 | 542434bp | 0.02% |
| –SVA: | 1955 | 540441bp | 0.02% | 1955 | 542434bp | 0.02% |
| **SINE:** | 321096 | 50874809bp | 1.59% | 321169 | 51519142bp | 1.61% |
| –5S: | 112 | 6669bp | 0.00% | 112 | 6575bp | 0.00% |
| –5S-Deu-L2: | 6 | 657bp | 0.00% | 6 | 657bp | 0.00% |
| –5S-Sauria-RTE: | 11 | 2942bp | 0.00% | 11 | 2942bp | 0.00% |
| –Alu: | 316135 | 49564845bp | 1.54% | 316208 | 50330316bp | 1.57% |
| –ID: | 101 | 7308bp | 0.00% | 101 | 7308bp | 0.00% |
| –MIR: | 4583 | 1638673bp | 0.05% | 4583 | 1638673bp | 0.05% |
| –U: | 82 | 25802bp | 0.00% | 82 | 25826bp | 0.00% |
| –tRNA: | 8 | 1428bp | 0.00% | 8 | 1428bp | 0.00% |
| –tRNA-Core-L2: | 16 | 2392bp | 0.00% | 16 | 2392bp | 0.00% |
| –tRNA-Core-RTE: | 13 | 680bp | 0.00% | 13 | 680bp | 0.00% |
| –tRNA-RTE: | 29 | 7788bp | 0.00% | 29 | 7788bp | 0.00% |
| **SINE?:** | 15 | 16871bp | 0.00% | 15 | 16871bp | 0.00% |
| –OTHER: | 15 | 16871bp | 0.00% | 15 | 16871bp | 0.00% |
| **Satellite:** | 21186 | 15224192bp | 0.47% | 21200 | 15239161bp | 0.47% |
| –OTHER: | 11836 | 886823bp | 0.03% | 11849 | 889852bp | 0.03% |
| –Y-chromosome: | 6796 | 13796614bp | 0.43% | 6796 | 13807838bp | 0.43% |
| –acromeric: | 655 | 173336bp | 0.01% | 655 | 173336bp | 0.01% |
| –centromeric: | 1421 | 322663bp | 0.01% | 1422 | 323379bp | 0.01% |
| –telomeric: | 478 | 65901bp | 0.00% | 478 | 65901bp | 0.00% |
| **Simple:** | 19907 | 915828bp | 0.03% | 19927 | 919915bp | 0.03% |
| –repeat: | 19907 | 915828bp | 0.03% | 19927 | 919915bp | 0.03% |
| **Unknown:** | 96437 | 19902744bp | 0.62% | 96738 | 20052302bp | 0.62% |
| –OTHER: | 96437 | 19902744bp | 0.62% | 96738 | 20052302bp | 0.62% |
| **rRNA:** | 586 | 73310bp | 0.00% | 586 | 73310bp | 0.00% |
| –OTHER: | 586 | 73310bp | 0.00% | 586 | 73310bp | 0.00% |
| **scRNA:** | 247 | 22162bp | 0.00% | 247 | 22162bp | 0.00% |
| –OTHER: | 247 | 22162bp | 0.00% | 247 | 22162bp | 0.00% |
| **snRNA:** | 329 | 24069bp | 0.00% | 329 | 24052bp | 0.00% |
| –OTHER: | 329 | 24069bp | 0.00% | 329 | 24052bp | 0.00% |
| **tRNA:** | 498 | 54792bp | 0.00% | 498 | 54792bp | 0.00% |
| –OTHER: | 498 | 54792bp | 0.00% | 498 | 54792bp | 0.00% |
